# A unified rheological model for cells and cellularised materials

**DOI:** 10.1101/543330

**Authors:** A Bonfanti, J Fouchard, N Khalilgharibi, G Charras, A Kabla

## Abstract

The mechanical response of single cells and tissues exhibits a broad distribution of time scales that gives often rise to a distinctive power-law regime. Such complex behaviour cannot be easily captured by traditional rheological approaches, making material characterisation and predictive modelling very challenging. Here, we present a novel model combining conventional viscoelastic elements with fractional calculus that successfully captures the macroscopic relaxation response of epithelial monolayers. The parameters extracted from the fitting of the relaxation modulus allow prediction of the response of the same material to slow stretch and creep, indicating that the model captured intrinsic material properties. Two characteristic times can be derived from the model parameters, and together these explain different qualitative behaviours observed in creep after genetic and chemical treatments. We compared the response of tissues with the behaviour of single cells as well as intra and extra-cellular components, and linked the power-law behaviour of the epithelium to the dynamics of the cell cortex. Such a unified model for the mechanical response of biological materials provides a novel and robust mathematical approach for diagnostic methods based on mechanical traits as well as more accurate computational models of tissues mechanics.

---

As part of their physiological function, single cells and tissues are continuously exposed to mechanical stress. For example, leukocytes circulating in the blood must squeeze through small capillaries, and the epidermis must deform in response to movements of our limbs. During development, mechanical forces initiate morphogenetic processes involving epithelial growth, elongation or bending, acting as cues to coordinate morphogenetic events [1]. Epithelial cell sheets are also continuously subjected to deformation as part of normal physiology. For instance, lung epithelial cells are exposed to fast cyclical mechanical stress during respiration [2], while epithelia lining the intestinal wall or those in the skin can experience long lasting strain [3]. Failure to withstand physiological forces results in fracture of monolayers which may lead to severe clinical conditions, such as hemorrhage or pressure ulcers [3]–[6]. Despite significant progress with the experimental characterization of cell and tissue mechanics, understanding the role of mechanical forces in development and pathology is hampered by the lack of a unified quantitative approach to capture, compare and predict the complex mechanical behaviours of tissues, cells, and sub-cellular components across all physiologically relevant time-scales. Such a framework would also enable us to assess the effects of pharmacological treatments on tissue mechanical response without necessitating experimental characterization of the tissue response to all loading conditions, something important for tissue engineering and the design of palliative treatment strategies.

In recent years, experimental characterization of the mechanical behaviour of single cells and tissues has revealed a complex set of mechanical behaviours in response to deformation [7]–[10]. For example, both single cells and tissues often display multiphasic responses in stress relaxation and creep tests, which comprise a combination of power-law and exponential behaviours. Power-law responses are commonly observed in biomaterials and are thought to originate from their complex hierarchical structure [11]–[14]. These behaviours cannot be easily modelled using traditional linear viscoelasticity, where constitutive rheological models result from combinations of elastic springs and viscous dashpots that translate into sets of linear ordinary differential equations [9], [15]–[18]. In this framework, power laws can only be implemented through a large numbers of linear elements [19], making this approach impractical and uninformative. Empirical functions have been introduced to overcome this challenge [11], [20], [21], but the lack of underlying material model, in the form of a computable differential equation, prevents the direct comparison of data collected under different loading conditions.

One potential approach for modelling the mechanics of materials presenting power-law behaviours is fractional calculus [22]. This relies on the introduction of a mechanical viscoelastic element called a spring-pot whose behaviour is intermediate between a spring and a dashpot [23]. This element based on fractional derivatives captures, with only two parameters, the broad distribution of characteristic times [24] typical of the mechanical response of cellularised materials. This element has recently been combined with traditional elements to model more complex rheological behaviours, referred to as generalized viscoelastic models [25].

In this paper, we examine the potential of generalized viscoelastic models for modelling biological materials by combining traditional rheological elements with the springpot. With only four parameters, we capture the time-dependent response of single cells and epithelial monolayers. Using parameters extracted from relaxation tests, we are able to predict the response of the same material to creep and ramp deformations with no further fitting, and relate the model parameters to single cell characteristics as well as recent measurements of cortical rheology.

## A constitutive model for epithelial monolayers

One widely used model system for studying tissue mechanics is the epithelium monolayer. We focused on MDCK cell monolayers devoid of substrate, of typical width of 2 mm and suspended between two rods at a distance of 1.5 mm. Despite the absence of a substrate, cells still retain epithelial characteristics [9]. Such a material has now been extensively studied [26], [27], and the effect of pharmacological treatments on rheological properties characterized [20]. The advantage of such a simplified system lies in the fact that the tension is transmitted only through the intercellular junctions and the cytoskeleton, but not the extracellular matrix. The relaxation response (response to a step in strain, see experimental details in section 1 in Supplementary materials) consists of an initial power-law phase in the first 5 s, followed by an exponential phase that reaches a plateau at ∼60 s (figure 1) [20]. This description establishes the minimum number of parameters needed to describe such a rich behaviour: (i) the level of the final plateau, (ii) the exponential decay time, (iii) the power-law exponent and (iv) the transition time between the two phases. This qualitative analysis will now inform the development of a novel rheological model tailored to capture these four components of the response using the minimum number of parameters.

**Figure 1:**
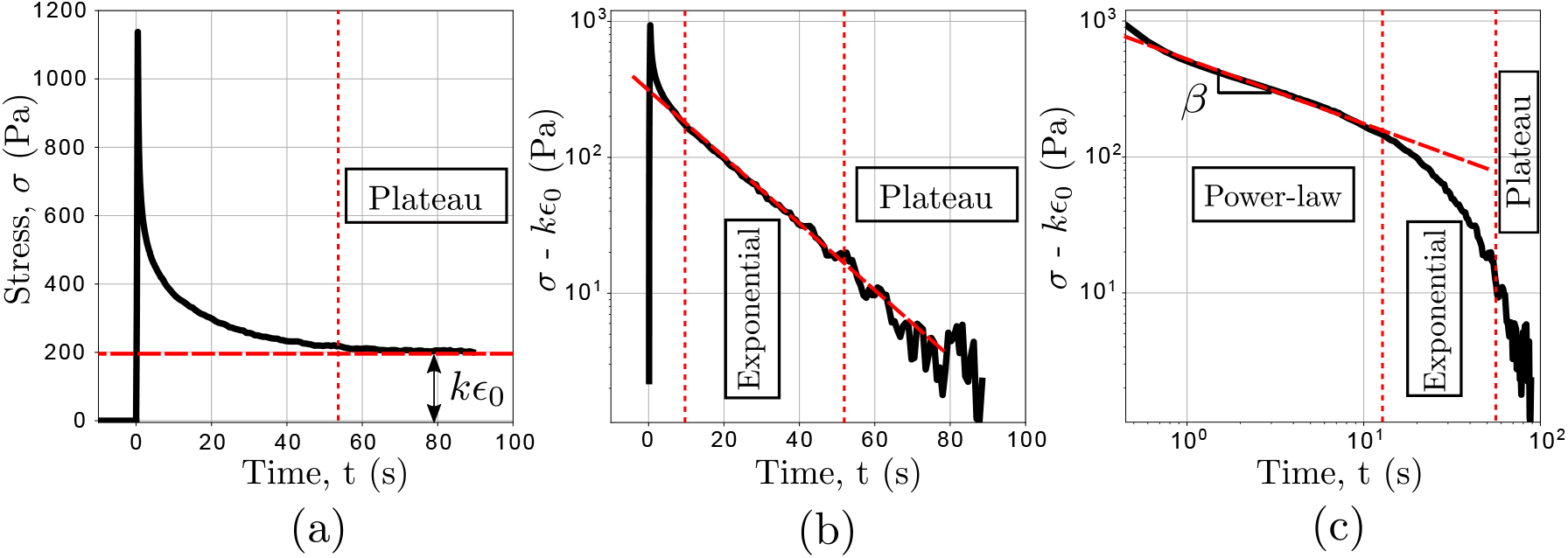
Representative experimental data of the stress relaxation of epithelial monolayers depleted of substrate. (previously reported in [20]). (a) An example of relaxation curve which highlights the final plateau. After removing the plateau, the relaxation curve is plotted in semilogarithmic scale (b) and logarithmic scale (c) to identify respectively the exponential and power-law behaviours.

A branch of Mathematics called Fractional Calculus provides conceptual and numerical tools well suited to capture power law behaviours [28], [29]. In traditional calculus, a function can be differentiated *n* times, where *n* is an integer. For viscous (fluid-like) materials, the stress is proportional to the first time derivative of the strain, where *n* = 1. For elastic (solid-like) materials, the stress is proportional to the strain, which can be seen as the zero-th time derivative of the strain *n* = 0. Fractional calculus generalizes the differentiation process such that the number *n* can now be real (see Supplementary materials section 2). With the spring-pot fractional element, the stress is proportional to the *β* derivative of the strain, where 0 ≤ *β* ≤ 1:

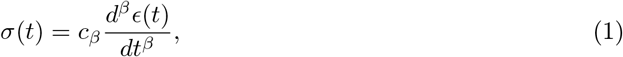

where *c*_*β*_ is a constant dependent on the material and *d*^*β*^/*dt*^*β*^ is the fractional derivative operator. When *β* = 0, the material behaves like a spring, and, when *β* = 1, like a dash-pot. As *β* varies from 0 to 1, the response of the material continuously transitions from elastic to viscous behaviour and if a step change in stress or strain is applied, the response exhibits a power-law. Mathematically, the response is only defined by an integral over time, leading to strong history dependence, referred to as the hereditary phenomena (Supplementary materials section 2). Despite this complexity, the spring-pot still lies in the linear viscoelastic framework, enabling us to greatly simplify the analysis of the data and make predictions. For dimensional consistency, the unit of the constant *c*_*β*_ is (Pa s^−*β*^), and therefore it does not have a straightforward physical meaning, although, it has been argued that it may represent a measurement of the firmness of the material [30].

The spring-pot can be combined with other rheological elements to generate a rich set of behaviours [31]. Configurations explored so far were mostly selected for their mathematical simplicity, rather than relevance to particular physical systems [25], [32]. Here, we adopt a phenomenological approach based on our qualitative description of the material’s behaviour, aiming to capture both its short and long time-scale response. At long time scale, the stress response shows a plateau (figure 1 (a)). Hence the model requires a spring in parallel with a dissipative branch that would not carry any tension in steady state. At intermediate time scales, the stress relaxes exponentially towards the plateau (figure 1 (b)). Hence the dissipative branch should behave as a dash-pot in series with an element that would transiently store deformation, for instance a spring as in a Maxwell model [33]. However, at short time scale (0.1-10 s), the dissipative branch exhibits a power-law. The presence of a power-law relaxation response immediately after application of strain, rather than a specific jump in stress, indicates that we should replace the spring in the dissipative branch with a spring-pot (see figure 2 (a)). The fractional model introduced in the dissipative branch is a special case of a known combination referred to as a Fractional Maxwell Model (FMM) [31]. The constitutive equation for the fractional material model introduced here in figure 2 (a) is reported in the Supplementary materials section 2. In what follows, such a qualitative approach is validated against experimental data, leading to a predictive model of the material’s behaviour.

**Figure 2:**
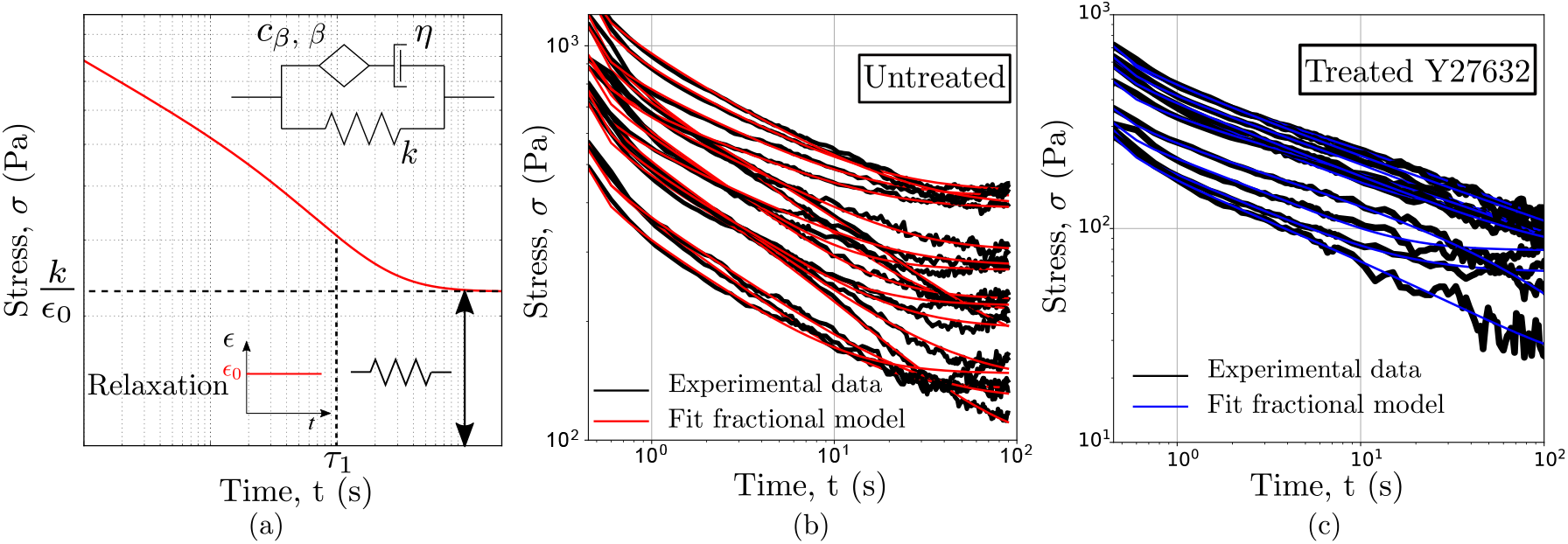
Fractional viscoelastic model for epithelial monolayers: constitutive model and stress relaxation behaviour. (a) Diagrammatic representation of the fractional rheological model and qualitative behaviour of its stress relaxation modulus. The spring combined with the spring-pot gives the initial response to a step in strain occurring at t=0 s. After the power-law decay, the curve converges to a plateau set by the stiffness of the spring. The characteristic time *τ*_1_dictates the transition between power-law and exponential regimes. The three-element fractional model is fitted to the relaxation data for (b) untreated epithelial monolayers (black curves are the experimental data, while the red curves represent the fit) and (c) monolayers treated with an inhibitor of contractility, Y27632 (the black curves are the experimental data, while the blue curves are the fits).

## The generalized fractional viscoelastic model characterizes the biphasic stress relaxation response

The relaxation modulus (stress response to a unit strain of deformation) of the fractional network model introduced above can be derived analytically. Since the relaxation modulus of two elements in parallel is given by the sum of their relaxation moduli, the relaxation modulus *G*(*t*) of the novel viscoelastic model presented in figure 2 (a) is obtained by adding the relaxation modulus of the Fractional Maxwell Model (FMM) to the stiffness of the spring, which results in

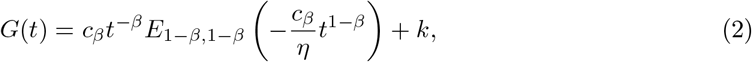

where *E*_*a,b*_ (*z*) is the Mittag-Leffler function, a special function that arises from the solution of fractional differential equations (see Supplementary materials section 2). The qualitative behaviour of the relaxation modulus is plotted in log-log scale in figure 2 (a). Since the argument of the Mittag-Leffler function is non-dimensional, we can identify a first characteristic time *τ*_1_, given by which approximates the transition time between the power-law and the exponential regime. The significance of *τ*_1_ as a characteristic relaxation time will be evident when the model is used to analyze real data.

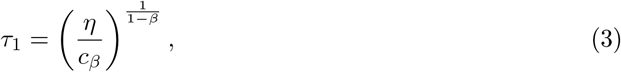

To assess the validity of the fractional model, we used it to fit the relaxation response of epithelial monolayers (see Supplementary materials section 1 for details of the experimental set up). In agreement with the qualitative analysis of the curves (figure 1), the four parameters involved in equation 2 account for the experimental data (see figure 2 (b)), successfully capturing all time domains: the power-law regime (for t < 10 s), the exponential behaviours and the steady-state stress. We further examined if the model could capture the relaxation response of epithelial sheets in which myosin contractility, one of the most important component controlling cellular mechanical properties, was inhibited [34]. We observed indeed that the same four parameters could fit well the experimental data (figure 2 (c)). This allows us to compare the parameters extracted from treated monolayers with their control (DMSO treated, figure S6 in Supplementary materials). The viscosity *η* of treated monolayers doubles compared to the DMSO treated monolayers (figure S6 (c) in Supplementary materials), in line with cell-scale findings suggesting that dynamic contraction of actin filaments increases cell fluidity [7]. By contrast, a reduction of the stiffness *k* was observed (figure S6 (d) in Supplementary materials), which suggests that acto-myosin contractility mainly plays a role in stress dissipation at long time-scale, consistent with the conclusions previously presented by [7], [20]. Other treatments have been applied to monolayers to examine the role of actin network organization and crosslinkers, without observing significant variations in the relaxation response of monolayers (see figures S6 and S7 in Supplementary materials).

## The generalised fractional viscoelastic model predicts the response to different loading conditions with no further fitting

The model relies on the assumption that the material behaves linearly. To identify the linear domain, we examined the stress response at different strain amplitudes ranging from 20% to 50%. We can observe that the material parameters are almost constant until roughly 30% (see figure S8 in Supplementary materials), which provides an upper bound of the linear domain where we expect the model to be valid. Within this range, we can assess the predictive power of our rheological description of epithelial monolayers. We extracted a distribution of parameters from the stress relaxation data and used them to estimate the response of the material to different forms of mechanical stimulation. Good agreement between predictions and experiments over a broad range of testing protocols would signify that our description represents a constitutive model whose parameters can be seen as material properties. We first consider the stress response to a strain ramp applied at constant strain rate (1%/s). The predicted response for the untreated monolayers is shown in figure 3 (a), with 95% confidence interval (see sections 3 and 5 Supplementary materials for details about prediction and statistical analysis, and see figure S5 (a) for Y27632 treated monolayers). The experimental results and the analytical predictions are in excellent agreement with no free parameters.

**Figure 3:**
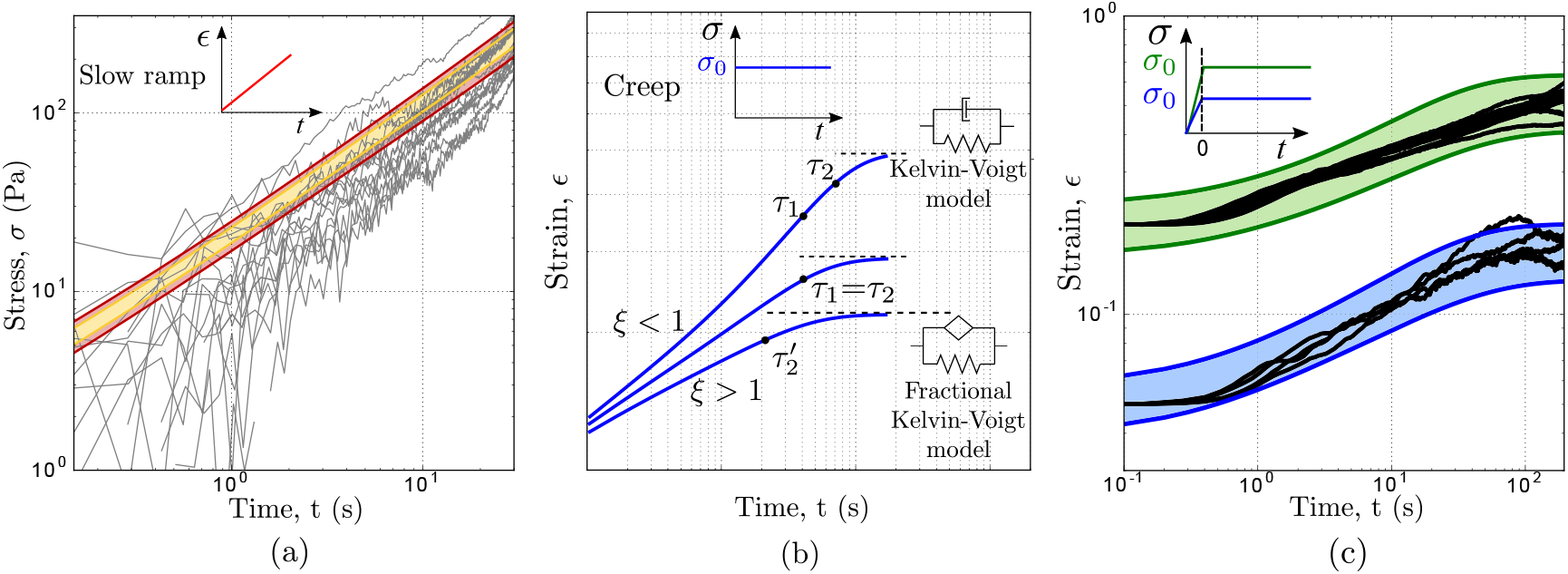
Prediction of epithelial monolayers response to different mechanical stimuli using the mechanical parametrization determined from stress relaxation experiments with no further fitting. (a) Predicted stress response of the untreated monolayers when subjected to a slow stretch (1%/s). The predicted responses (95% confidence interval red areas and 70% yellow areas) are in good agreement with the experimental data (black curves). The upper and lower limits of the predicted response are obtained by considering the standard error for each mechanical parameter. (b) The creep response of epithelial monolayers: fractional model response and experimental data. Sketch of the creep compliance of the generalized fractional viscoelastic model. Three possible qualitative behaviours can arise dependent on the relative values of the two characteristic times. (c) Creep response of the untreated epithelial monolayers. Two loadings are tested, 170 Pa (blue area) and 470 Pa (green area). These loads correspond to an initial strain respectively of 5% and 20%; therefore, the linearity assumption still holds. Note that the initial response of the creep is different from (b). This is due to the ramp during the initial phase (see section 1 Supplementary materials).

Similarly, we can challenge the model by predicting and validating the deformation response *J* (*t*) of the epithelial monolayers to a unit step in stress, a test usually referred to as a creep experiment (see section 3 Supplementary materials). For linear viscoelastic materials the relation between relaxation 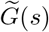 and creep 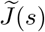 moduli in the Laplace domain is relatively simple, and given by 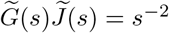 After transforming the relaxation modulus in equation (2) in the Laplace domain, we find:

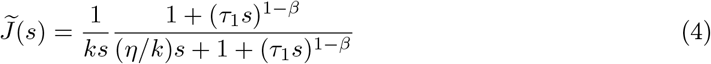

To obtain the solution in the time domain *J* (*t*), the inverse Laplace transform of the equation above is performed numerically.

The creep response is richer than the relaxation response, for which *k* only added a simple offset to the stress. Here, because the imposed load can continuously redistribute between the two branches of the model, *k* is involved in the dynamics. We can indeed identify in the creep response an additional time-scale *τ*_2_involved in the response: 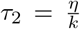 We therefore have one additional dimensionless parameter which controls the shape of the creep response *ξ τ*_1_,/*τ*_2_, The value of *ξ* leads to qualitatively different responses as plotted in figure 3 (b). (i) If *ξ* < 1, at short times we first observe a power-law behaviour arising from the spring-pot followed by an exponential regime where the dashpot dominates. The transition from spring-pot-dominated to dashpot-dominated regime is governed by the characteristic time *τ*_1_, as for the relaxation response. While the deformation increases, the spring eventually becomes relevant and the system tends towards the plateau as a Kelvin-Voigt model with a characteristic time *τ*_2_. (ii) If *ξ* > 1, the spring saturates before the transition to the dashpot occurs in the dissipative branch. Hence, the model behaves as a Fractional Kelvin-Voigt model with a characteristic time 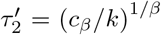 which can be expressed as 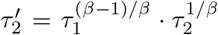is irrelevant in this regime. (iii) If *ξ* ≈ 1 the transition from the spring-pot to the dashpot corresponds to the time at which the spring becomes relevant.Therefore, the transition from the spring-pot to the Kelvin-Voigt model occurs with characteristic time *τ*_1_ ≈ *τ*_2_.

The model is now used to predict the response of monolayers when subjected to a step in stress using the material parameters derived from the relaxation experiments. We performed new experiments to test our model’s predictive power (see section 1 in Supplementary materials). In these experiments, we subjected monolayers to a stress which was maintained constant after an initial short ramp in strain (see inset figure 3 (c)). Strikingly, the experimental data falls well within the 95% confidence interval of the predicted response with no free parameters (details in section 3 and 5 in Supplementary materials) (figure 3 (c)).

MDCK monolayers seem to exhibit very different creep behaviour when myosin II activity is reduced. Whilst untreated monolayers reach a steady strain value after about 100 s, Y-27632 treated tissues continue to flow in a power-law manner (see figure S5), suggesting a qualitative difference between the two systems. This apparent contrast is however properly accounted for by the model. The predicted creep responses, calculated using the parameters obtained from figure 2, are in good agreement with the experimental data with no free parameters. Based on the parameters obtained form relaxation experiments on Y-27632 treated monolayers, *τ*_1_ and *τ*_2_ are both much larger than in the untreated case, and now comparable in value with the duration of the measurement (see table S2 in Supplementary materials). Furthermore, the reduction of *ξ* in the treated case, down to 0.7 compared to about 1 in the untreated case (figure S5 (c)), would change the shape of creep curve and give the impression that creep accelerates rather than saturates at times close to *τ*_1_(figure 3(b)). This analysis illustrates how modelling can bring consistency across systems that may at first appear qualitatively different, in particular when observed over a finite experimental time.

## Usage of the model beyond epithelial monolayers

Building on the work on MDCK monolayers, we may now consistently analyze data across biological systems, and pull information from different research groups, working with different mechanical testing protocols. For instance, we can show that the model successfully captures the relaxation response of single isolated cells, such as epithelial MDCK cells (figure 4 (a) [20], details on experimental setup in section 6) and articular chondrocytes (figure 4 (b), original data presented by Darling et. al. [35]).

**Figure 4:**
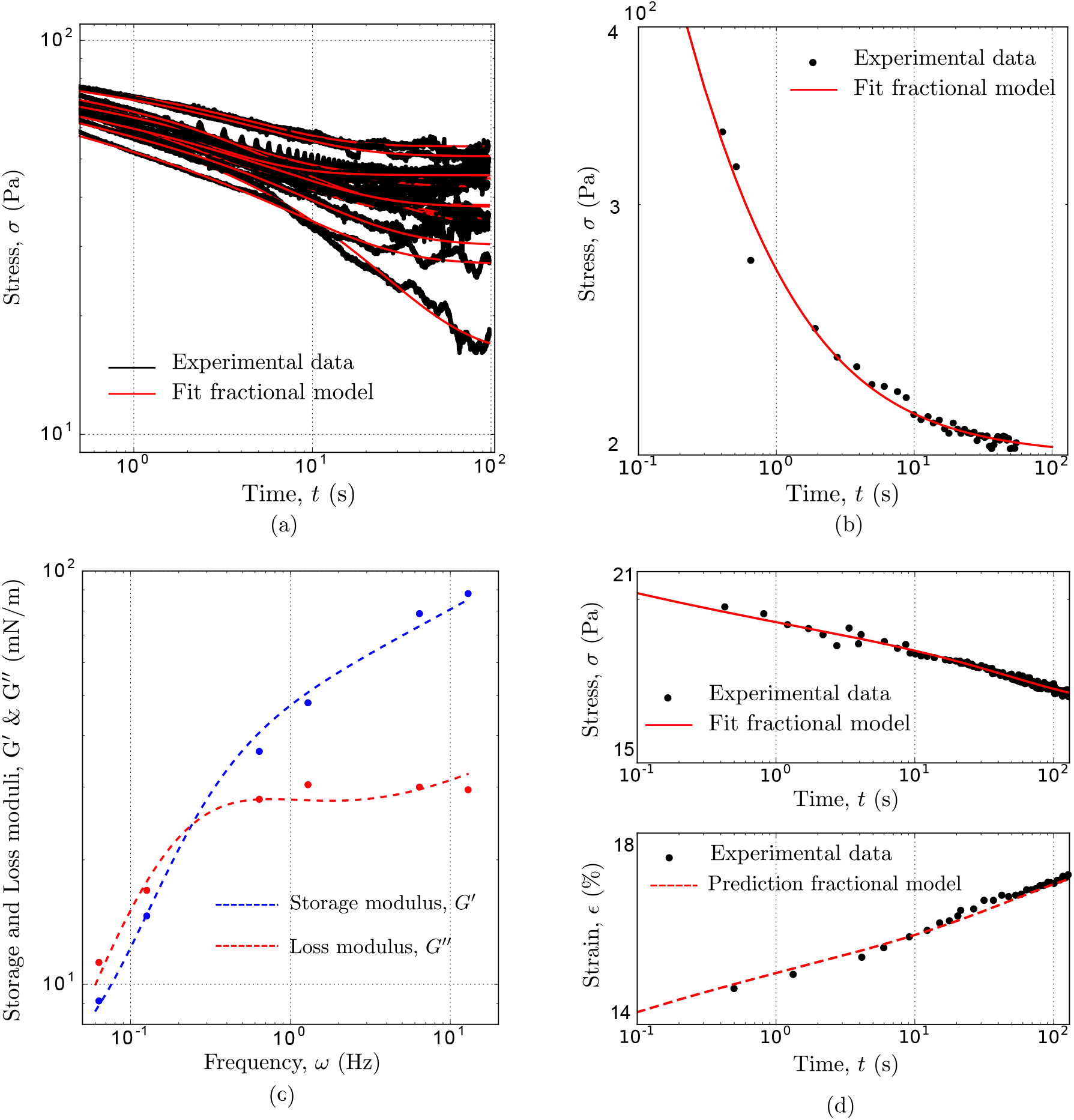
The unified model captures the mechanical behaviour regardless of the experimental set-up, the type of material and the length-scale. (a) Fitting of the relaxation response of single epithelial cells presented by Khalilgharibi et.al [20]. Each line is a different cell.(b) Fitting of the relaxation response of articular zone chondrocytes (original data from [35]) (c) Fitting of the storage and loss modulus of HeLa Kyoto cells (original data from [7]) using the model presented here. The blue and red dashed lines are respectively the fitted storage and loss modulus while the dots are the experimental data. (d) Fitting of the relaxation response of collagen fibrils (top graph) and prediction of the creep response (bottom graph). Original data from [36]. Fitted parameters in Table S2 in Supplementary materials.

Looking at sub-cellular components, the main factors controlling the mechanical properties of cells and tissues have been characterized experimentally already. Fischer-Friedrich *et al*. [7] have recently performed oscillatory compressions of HeLa Kyoto cells during mitosis by using an AFM cantilever and analysed the data to extract the rheological behaviour of the cell’s cortical actin network. To allow the direct comparison between our constitutive model and the rheological data introduced by Fischer-Friedrich *et al*. [7], we calculated the analytical expression of the complex modulus *G*′ (*ω*)+ i*G*″(*ω*) associated with our model (see section 7 in Supplementary materials) and fitted the experimental data again with an excellent agreement as shown in figure 4 (c).

Likewise, with the modelling framework presented here we have been able to capture the relaxation response of collagen fibrils (figure 4 (d)-top). Furthermore, using the parameters extracted from fitting the relaxation data, we have been able to predict their creep behaviour (figure 4 (d)-bottom); a quantitative link that was absent in the original paper presented by Shen *et al*. [36].

As many biomaterials exhibit power law rheology, examples where the generalized fractional viscoelastic model successfully captures their behaviour abound in literature–e.g. blastomere cytoplasm and yolk cell rheology (see figure S12 (c)-(d) in Supplementary materials). Studies available in the literature have reported simpler qualitative creep and relaxation behaviour for single cells, such as a single power-law or a two-power-law behaviour [37]. These responses are embodied as special cases of the presented generalized model (negligible stiffness and/or large viscosity). To illustrate this, we fitted the power-law response of single immune cells (figure S13). We could also capture the creep response of a single muscle cell exhibiting a two-power-law behaviour, and predict its storage and loss modulus (figure S12 (a)-(b)). This sets the fractional viscoelastic model presented here as a promising tool to support a unified description of the mechanical response of a broad variety of biological tissues.

## Links with biophysical analysis

Fractional models capture with a few parameters complex power law behaviours commonly associated with a broad distribution of relaxation times. A single spring-pot element is for instance sufficient to model a power law response, with two parameters. The complexity and richness of the spring-pot is apparent in the way the fractional derivatives are calculated, requiring an integration over the whole deformation history of the material, whereas the response of traditional elements only depend on the instantaneous values or rate of change of the strain. This integral expression of the derivative is responsible for the strong history dependent effects associated with power law rheology. The approach becomes even more powerful when combining the spring-pot element with other rheological components. Based on dimensional analysis, we identified two characteristic times and an effective stiffness involved in master curves for both the relaxation and creep functions. In the case of creep, the ratio of the two time-scales controls the qualitative shape of the response, accounting for seemingly different responses of the material with and without contractility. This provides ways to bridge our mathematical framework with more physical descriptions of the rheology of cells and tissues. For example, Fischer-Friedrich *et al*. [7], after observing that a simple power law function failed to capture the dynamic response of HeLa Kyoto cells, built a physical model based on a flat distribution of relaxation times up to a certain cut-off time. The model we present here captures a similar phenomenology by introducing a dashpot in series with a springpot. The characteristic time *τ*_1_defined for the fractional model acts as a cut-off in the relaxation spectrum, causing a transition from a power-law to an exponential behaviour in the relaxation function. In both studies, a reduction of myosin activity leads to an increase of the characteristic or cut-off time scale. The model proved to be useful to compare two different biological systems, Hela and MDCK, and identify similar behaviours despite the use of distinct testing approaches.

More generally, having a unified language to consistently capture the response of materials over a broad range of time scales is a significant step towards the understanding of the physical and biological significance of rheological behaviours, one of the current key challenges in Mechanobiology [38]. For instance, by comparing the relaxation response of single MDCK cells with MDCK monolayers, we noticed that they display similar behaviours (figure 4 (a)). However, looking at the parameter values (see table S2) reveals that monolayers exhibit a higher stiffness *k*, “firmness” *c*_*β*,_ and viscosity *η*. Quantifying these mechanical properties raises novel questions, and we can only speculate at this stage about the reasons. The increment in stiffness *k* displayed by the suspended monolayers compared to single cells may for instance be ascribed to an increase in cortical contractility as previously reported [20]. By contrast, the origin of the higher “firmness” of the spring-pot and viscosity *η* could be related to changes in the internal cell organization when cells form intercellular junctions to integrate into a monolayer. In the context of single cells, the relaxation plateau observed when cells are deformed could be associated with cortical tension [7], confirming further a strong link between *k* and cortical activity.

Although cells often exhibit non-linear behaviours, the model provides a rich base line from which to assess unusual traits of their response. Recently, Bonakdar *et al*. [21] performed local measurements of the rheology of mouse embryonic fibroblasts by imposing cycles of loading at constant force and relaxation using a magnetic particle linked to the cytoskeleton through the cell membrane by a fibronectin coating (figure 5 (a)). They observed a power law response coupled with an incomplete cell recovery suggesting plastic deformations that tend to decrease in amplitude with the number of loading cycles. Such a local mode of deformation is difficult to model quantitatively due to the complex and undetermined interaction between the bead and cytoskeleton. We nonetheless simulated such cycles of loading for an arbitrary amplitude of imposed stress, using the model parameters previously obtained from the MDCK cells. At first, we only considered a simple power-law behaviour (i.e. a single spring-pot) and confirmed the results of Bonakdar *et al*. that a power-law alone cannot reproduce the progressive drift of the bead displacement or cell deformation observed experimentally (figure 5 (b)). We then tested the full model obtained for MDCK cells; here again, the model provides deformation that quickly return to 0, missing the deformation drift visible in the experimental data (figure 5 (c)). However, as discussed earlier, the spring *k* appears to be primarily linked to the overall deformation of the cell cortex, providing a restoring force if the cell is squashed or elongated. Given the local nature of Bonakdar *et al*.’s measurements, we expect that the elastic term *k* originating from cortical tension is likely to be irrelevant as a first approximation. By removing accordingly the contribution of the spring, the response to cycles of loading does provide a drift that is more consistent with experimental data, albeit growing linearly with time and exceeding the experimental trend (figure 5 (d)). However, it has been shown previously that during application of a local stress through fibronectin coated beads, cells respond with a local strengthening of the cytoskeleton linkages [39]. By reinstating a small proportion of the spring value to account for such local stiffening we could generate a series of curves consistent with the experimental data over the first 80 seconds. Although this analysis may not fully account for the non-linear plastic behaviour reported in mouse embryonic fibroblasts, it illustrates very clearly that a number of qualitative distinctive traits, reported in different systems and under different experimental conditions, may be in fact largely consistent with each other.

**Figure 5:**
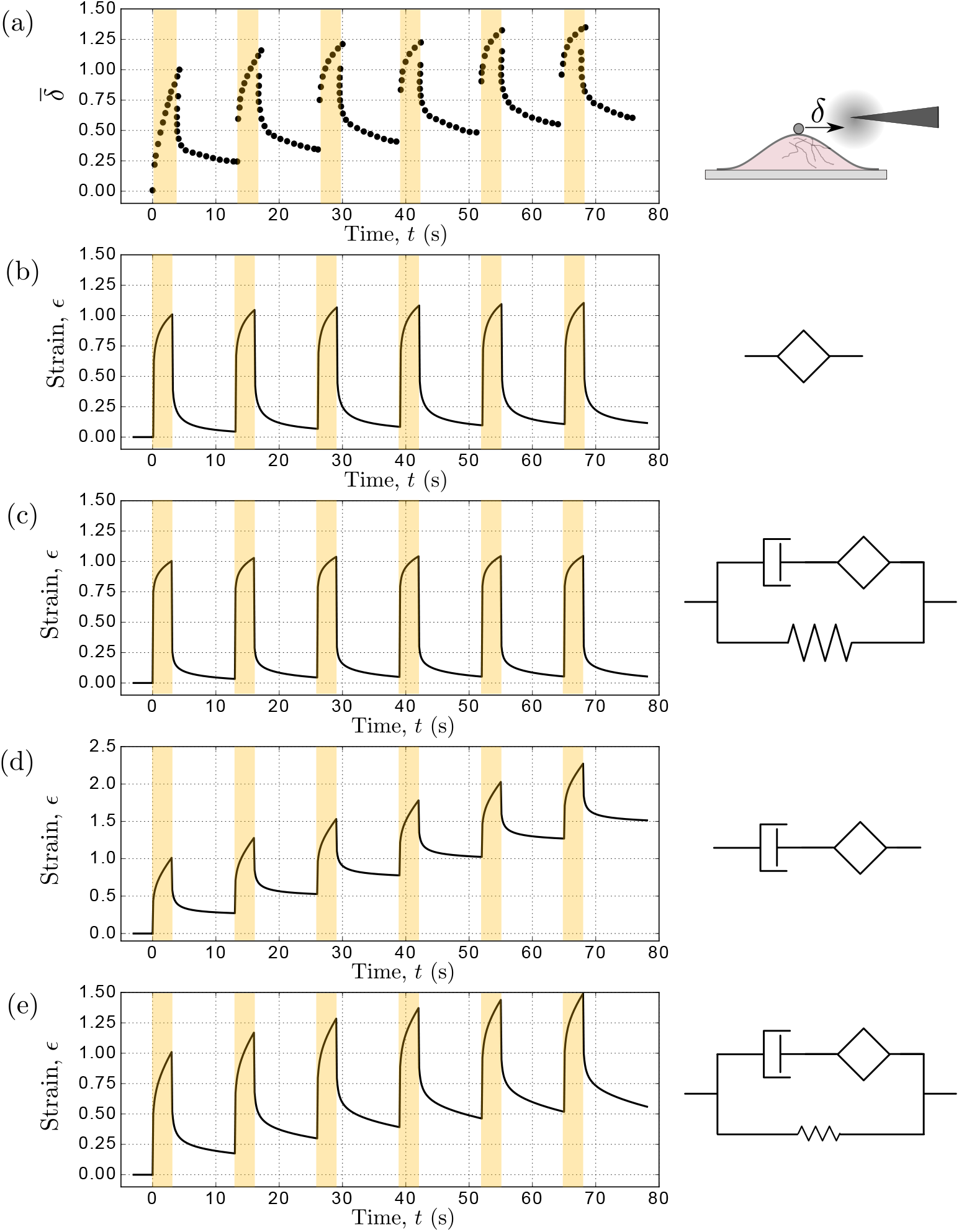
Analysis of local rheological data. (a) Incomplete recovery of a single cell deformation to alternating force cycles. Digitized data from Bonakdar *et al*. [21]. A magnetic bead is attached to the cytoskeleton via integrin-type adhesion receptors through which a force parallel to the cell is applied (force on in dark areas). Note that the displacement is normalized respect to the first peak value because it is difficult to identify the interaction of the bead with the cell and therefore convert the displacement into strain. By using the material parameters of MDCK cells (table S2), we predict the response to cycles of loading using the fractional models and we compare the qualitative responses normalized respect to the first peak (b)-(e) with the experimental data (a). Prediction of the response to cyclic loading with the spring-pot only (b), with the novel model (c), without cell contractility (thus *k* = 0) and with a small cell contractility (thus *k* = 20 Pa, 10% of the fitted value of *k*).

## Conclusions

We have presented a constitutive model for epithelial monolayers that captures the biphasic nature of their stress relaxation dynamics over their full physiological functional range. From a qualitative analysis of the monolayers’ relaxation response, we determined the number of parameters required to account for the behaviour observed, and combined traditional viscoelastic elements with the fractional spring-pot to fit tissue responses. The particular rheological model introduced here has been validated for the first time against experimental data for the relaxation response of suspended MDCK monolayers. Strikingly, the values for the parameters obtained by fitting the relaxation response could predict the monolayer’s response to other mechanical tests with excellent accuracy. This confirms that our model provides a constitutive description of the material and that the fitting parameters are proper material properties independent of the type of deformation or force applied to the material. We further demonstrated the model’s suitability to analyse and compare rheological behaviour across many systems and length scales, within the same unifying framework. It greatly facilitates a biophysical analysis by enabling a more intuitive approach to power-law modelling, and allowing us to ask more direct questions regarding the biological significance of the parameters involved.

Beyond enhancing our understanding of biological systems, a unified rheological model for biomaterials is also crucial in the medical and engineering fields. Correlating changes in the mechanical response of tissues to their biological state has been long considered as a promising marker-free method for cancer diagnosis [40]–[42]. In the context of regenerative medicine, rheological phenotypes provide a suitable metric to assess the similarity between tissue engineered constructs produced *in vitro* and their natural counterparts. As the parameters of the model are material properties that are independent of the method of measurement, they are promising mechanical signatures of the tissue and its condition to be used for diagnosis or as a target for regenerative medicine.

A practical limiting factor for the widespread application of the model presented here is the mathematical complexity of fractional derivatives, and the current lack of user-friendly numerical methods to perform such analysis without expertise in fractional calculus. Such tools have been recently released in the public domain by our group [43]. This will ensure a broader adoption of fractional models for a rapid and systematic analysis of experimental data, as well as future integration within numerical packages and finite element software.

## Acknowledgements

The authors wish to acknowledge present and past members of the Kabla and Charras labs for stimulating discussions. A.B. and A.K. wish to acknowledge Louis Kaplan for his important contribution to the development of the RHEOS package and Arran Fernandez for stimulating discussions on fractional calculus. A.B and J.F. were funded by BBSRC grant (BB/M003280 and BB/M002578) to G.C. and A.K. N.K. was part of the EPSRC funded doctoral training program CoMPLEX. N.K. was also funded by the Rosetrees Trust, and the UCL Graduate School. G.C. is supported by a consolidator grant from the European Research Council (MolCellTissMech, agreement 647186). The work was supported by BBSRC grants (BB/K018175/1, BB/M003280 and BB/M002578).

## Author contributions

A.B and A.K. designed the rheological model. N.K carried out the relaxation and ramp experiments. J.F. and G.C designed the creep experiments. J.F. carried out the creep experiments. A.B and A.K. performed data analysis. J.F. and G.C. contributed to physical interpretation of the data. All authors discussed the results and manuscript.

## Supplementary materials of “A unified rheological model for cells and cellularised materials”

### 1 Mechanical testing of suspended epithelial monolayers

Experiments were performed on suspended MDCK II epithelial monolayers generated as described in [1]. Cells stably expressing E-Cadherin-GFP were used. After digestion of collagen substrate, monolayers were tested at room temperature in Leibovitz’s L15 medium (Gibco, ThermoFisher) supplemented with 10% FBS (Sigma).

Relaxation experiments were performed as described in [2]. Briefly, a mechanical testing setup assembled on top of an inverted microscope (Olympus IX-71) was used. First, the Petri dish containing the stretcher device was secured to the microscope stage. Next, an opto-mechanical force transducer (SI-KG7A, WPI) with a tweezer-shaped mounting hook (SI-TM5-KG7A-97902, WPI) was mounted on a motorized translation stage (M126-DG1, Physike Instrumente). The tip of the mounting hook was then brought into contact with the flexible arm of the stretcher device. The monolayer, suspended between the two arms of the device, was stretched by moving the flexible arm away from the rigid arm, using the motorized stage. Monolayers were extended to 30% strain at a rate of 75%/s. The deformation was then maintained constant during stress relaxation. The force applied to the monolayer was calculated by subtracting the force applied by the flexible arm from the total measured force. Ramp experiments were conducted on the same setup and monolayers were extended at a rate of 1%/s. Force measurements in both tests was acquired at 6.7 Hz.

For creep experiments, the setup was slightly adapted (figure S1). The deformation was controlled by moving the rigid arm connected to one extremity of the monolayer while the force was measured on the other static arm. Monolayers were prestretched by 10% prior to experiments and left to rest for 5 minutes before starting testing. Then, an initial stretch step was applied at 1.5 mm/s (about 100%/s) and the force reached was clamped using a feedback algorithm (Labview, National Instruments) and updated at 7.5 Hz.

To monitor the shape of the tissue and ensure its integrity during the tests, bright-field images were taken at a rate of 1 image/s using a 2X objective (2X PLN, Olympus) and a CCD camera (PointGrey, Grasshopper 3).

### 2 The fractional element: springpot

The spring-pot rheological element exploits the mathematical concept of fractional derivative to model behaviours that are intermediate between a spring and a dashpot (see figure S2). Different definitions of fractional differentiation are available. Here we use the Caputo’s derivative definition; it has shown a better applicability to real problems, where initial conditions are known in terms of derivative of integer index [1]. If we assume that the system is at rest for time *t* ≤0, the fractional derivative is given by [2]

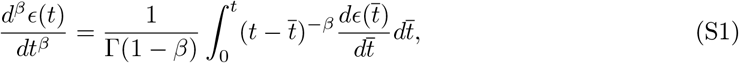

where Г(·) is the Gamma function and 0 < *β* < 1. Note that in order to get the value of the derivative at a given time *t*, it is necessary to integrate from *t* = 0. Thus, the response of the springpot is determined by its whole deformation history. This highlights the hereditary feature of such an element.

From a graphical representation of the linear viscoelastic model as a network of springs, dashpots and springpots, we can establish the time-dependent differential equation describing the relation between stress and strain. For the model presented here the temporal evolution is given by [3]

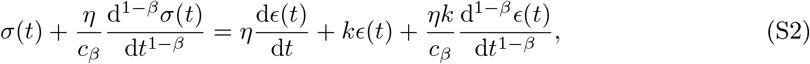

where *η* is the viscosity of the dashpot, *c*_*β*_ and *β* are the parameters of the spring-pot and *k* the stiffness of the spring.

**Figure S1:**
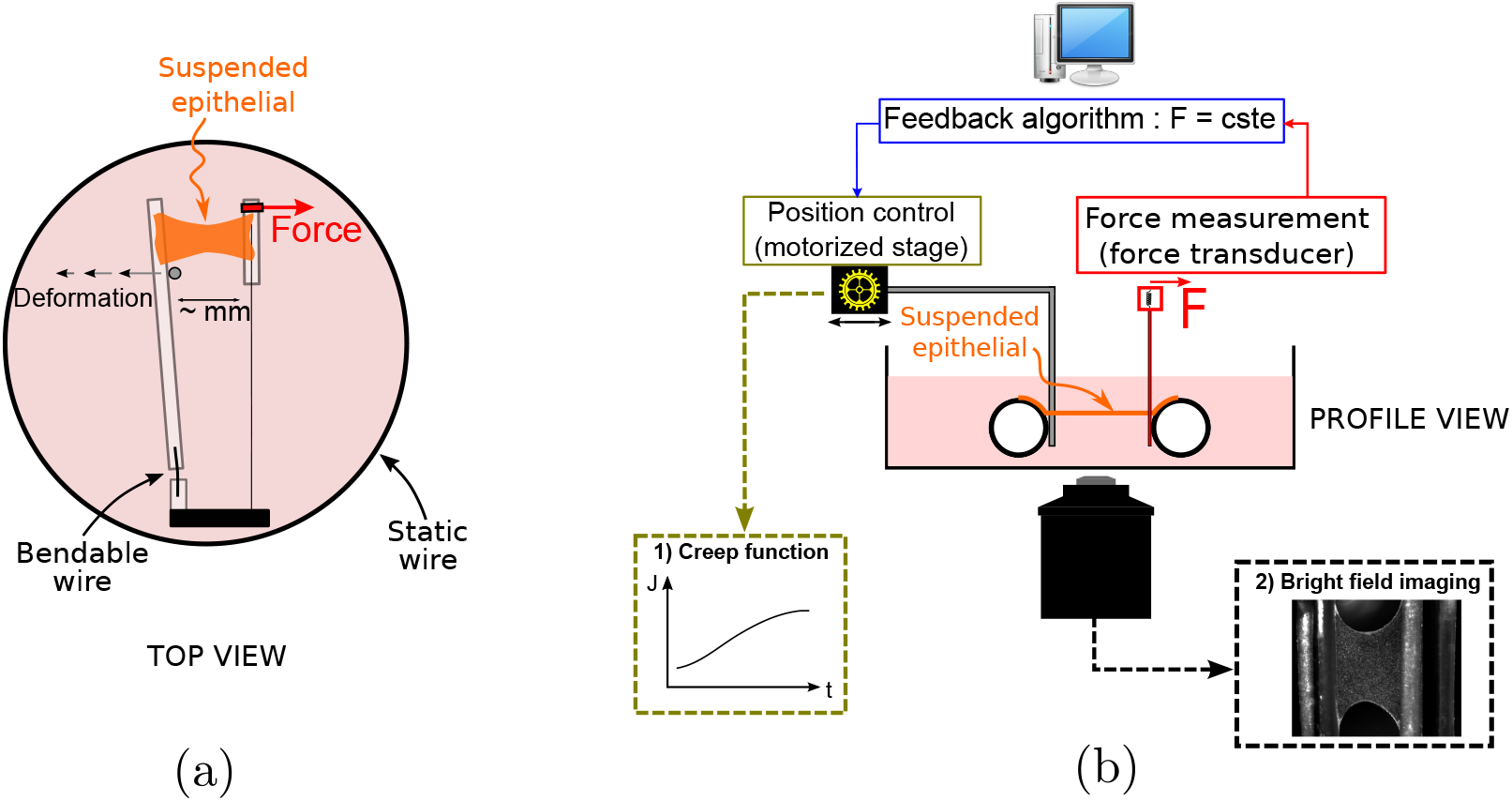
Set up of the creep experiment. (a) A millimetric MDCK epithelial monolayer devoid of substrate is suspended between two test rods. The device is glued to the Petri dish. The force is monitored on one side of the tissue, while the deformation is imposed on the other side. (b) A constant stress in the tissue is maintained by monitoring the traction force on the tissue through a force transducer (right) and adjusting its length in real time via a motorized stage. A PID feedback algorithm ensures the control of the stress

Many solutions of fractional differential equations obtained based on generalized viscoelastic models admit a closed form involving the Mittag-Leffler function [3]. Such a function is defined as

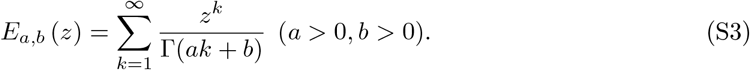

Its qualitative behaviour for the parameters of the soft material analyzed here is plotted in figure S3.

### 3 Curve fitting and prediction

Relaxation curves collected during stress relaxation tests were analyzed using the open source library RHEOS [4] developed in *Julia* language [5].

During a typical experimental test, the strain *E*(*t*) is rapidly applied at a constant rate until it reaches 30% and then maintained constant. The stress is obtain from the Boltzman integral, given by the convolution between the relaxation modulus and the derivative of the strain

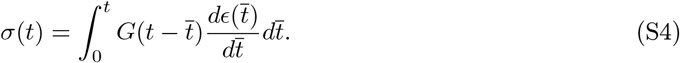

where *G*(*t*) is the relaxation modulus which takes the form reported in equation (2) in the manuscript.

The viscoelastic model presented here is consistent with the empirical function introduced in [6]

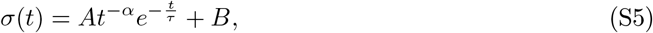

where *B* is the residual stress, *α* is the power exponent and *τ* is the characteristic time. To demonstrate this, we first re-fited the relaxation response using the same fitting procedure as in [6]. Thus *k* is fixed at the residual stress after reaching the plateau defined as the average of the stress for 70 s < *t* < 75 s. The parameters obtained for the viscoelastic model and the empirical expression are similar (see table S1). If *k* is left free during the fitting, a variation in the relaxation characteristic time is noticed between the two approaches, especially in the Y27632 parameters.

**Figure S2:**
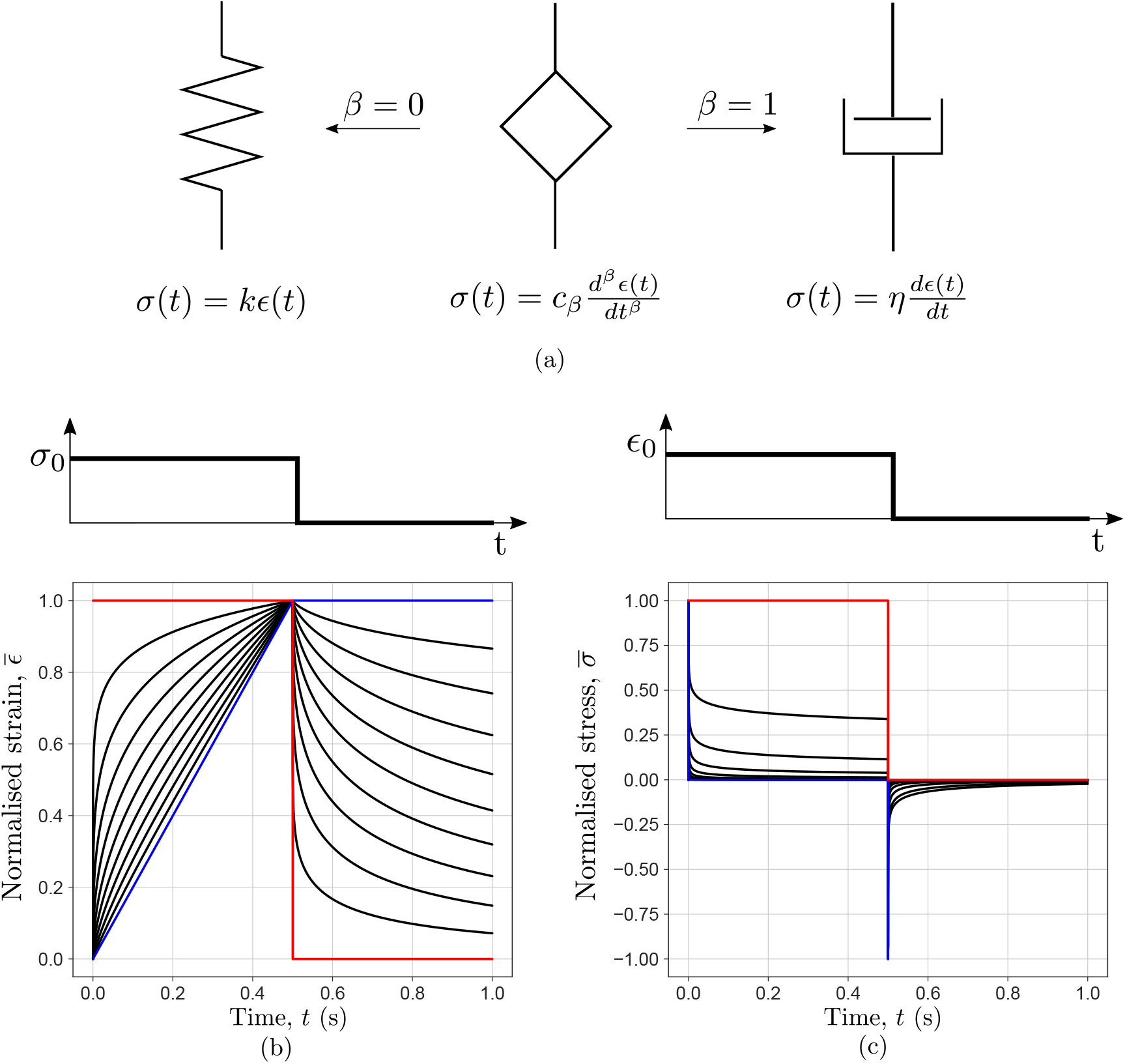
(a) Sketch of the fractional element–spring-pot. The spring-pot interpolates between a spring and a dashpot. When *β* = 0, the spring-pot reduces to a spring; whilst when *β* = 1 it becomes a dashpot. Consequently, the material constant *c*_*β*_ represents respectively the elastic constant of a spring, *k* (Pa) and the viscosity of a dashpot, *η* (Pa s). (b) Normalized responses of a springpot when subjected to a step in stress and (c) to a step in strain. The red curves are for a spring (*β* = 0), the black curves are for increasing values of *β* from 0.1 to 0.9 and the blue curves are for a dashpot (*β* = 1).

This discrepancy could be associated to the lack of a clear plateau in the experimental time-window. For the analysis presented in this work, we have used those parameters obtained by letting *k* free during the fitting.

To predict the response to a slow ramp, the Boltzman integral reported in equation S4 is used. The relaxation modulus *G*(*t*) takes the form reported in equation (2) in the manuscript, while the derivative of the strain will be equal to the applied strain rate. Note that the parameters of the relaxation modulus are those predicted from the fitting of the relaxation experiments.

The prediction of the creep response is obtained by solving the Boltzman integral

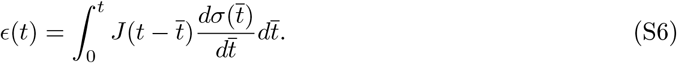

where *J* (*t*) is the creep modulus. To calculate the creep modulus we use the same parameters as extracted from the fitting of the relaxation experiments. The predicted creep response for the treated monolayers Y27632 is reported in figure S5 in the Supplementary materials.

**Figure S3:**
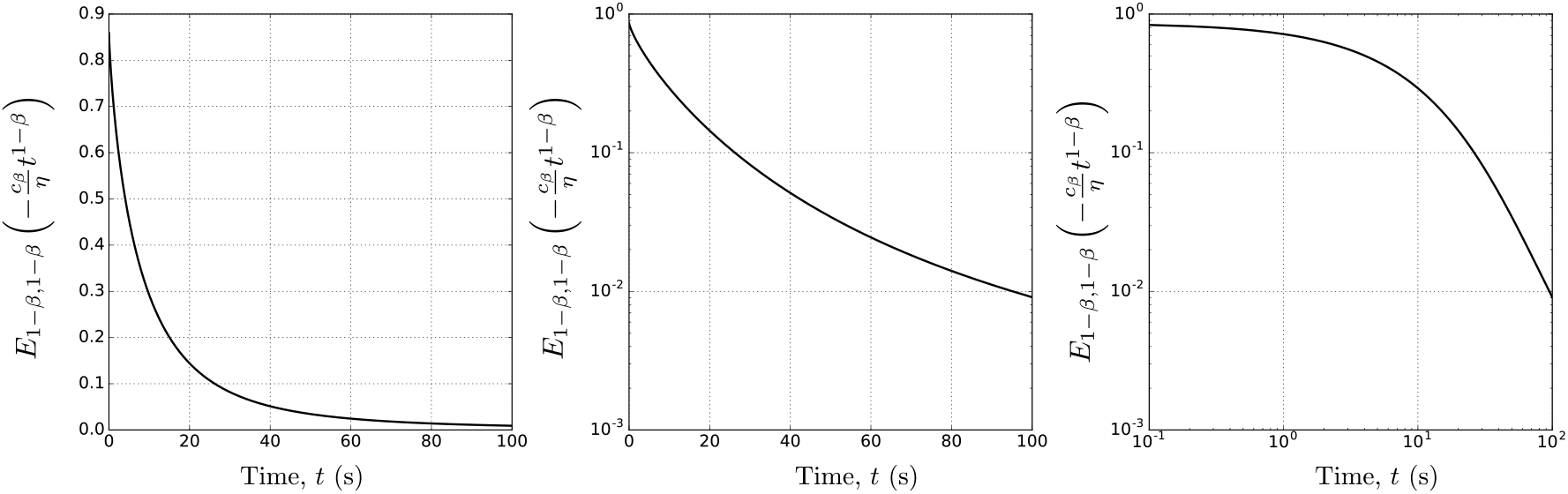
Mittag-Leffler function. Plot of the Mittag-Leffler function with parameters similar to those obtained for epithelial sheets (*β* = 0.22, *c*_*β*_ = 1.3× 10^3^ Pa s^*β*^, *η* = 1.0 × 10^4^ Pa s) in (a) linear, (b) semilogarithmic and (c) logarithmic scales.

**Table S1:** Comparison within the parameters *β* power exponent, *k* stiffness of the spring and *τ* characteristic relaxation time obtained from the fitting of: (a) the empirical function in [6], (b) the model presented here when *k* is fixed, as done in [6] and (c) when *k* is free.

| Sample |  | Control | DMSO | Y27362 | CK666 | SMIFH2 |
| --- | --- | --- | --- | --- | --- | --- |
| $\beta$ | Empirical | $0.28 \pm 0.02$ | $0.29 \pm 0.03$ | $0.26 \pm 0.05$ | $0.31 \pm 0.01$ | $0.29 \pm 0.02$ |
| | $k$ fixed | $0.30 \pm 0.03$ | $0.27 \pm 0.02$ | $0.287 \pm 0.01$ | $0.29 \pm 0.02$ | $0.28 \pm 0.02$ |
| | $k$ free | $0.28 \pm 0.03$ | $0.27 \pm 0.02$ | $0.296 \pm 0.05$ | $0.30 \pm 0.02$ | $0.29 \pm 0.03$ |
| $k$ (Pa) | Empirical | $820 \pm 373$ | $1130 \pm 327$ | $303 \pm 97$ | $1290 \pm 340$ | $1213 \pm 323$ |
| | $k$ fixed | $850 \pm 364$ | $1270 \pm 492$ | $300 \pm 101$ | $1286 \pm 340$ | $1212 \pm 321$ |
| | $k$ free | $840 \pm 376$ | $1200 \pm 428$ | $187 \pm 101$ | $1290 \pm 337$ | $1194 \pm 348$ |
| $\tau$ (s) | Empirical | $11.3 \pm 3.8$ | $13.3 \pm 5.5$ | $23.3 \pm 7.4$ | $16.8 \pm 6.2$ | $19.5 \pm 3.5$ |
| | $k$ fixed | $17.2 \pm 7.1$ | $11.8 \pm 4.31$ | $24.2 \pm 12.7$ | $20.42 \pm 13$ | $21.2 \pm 11.5$ |
| | $k$ free | $20 \pm 15.2$ | $13.58 \pm 11.6$ | $213.83 \pm 246$ | $19.51 \pm 14$ | $34.27 \pm 34$ |

**Figure S4:**
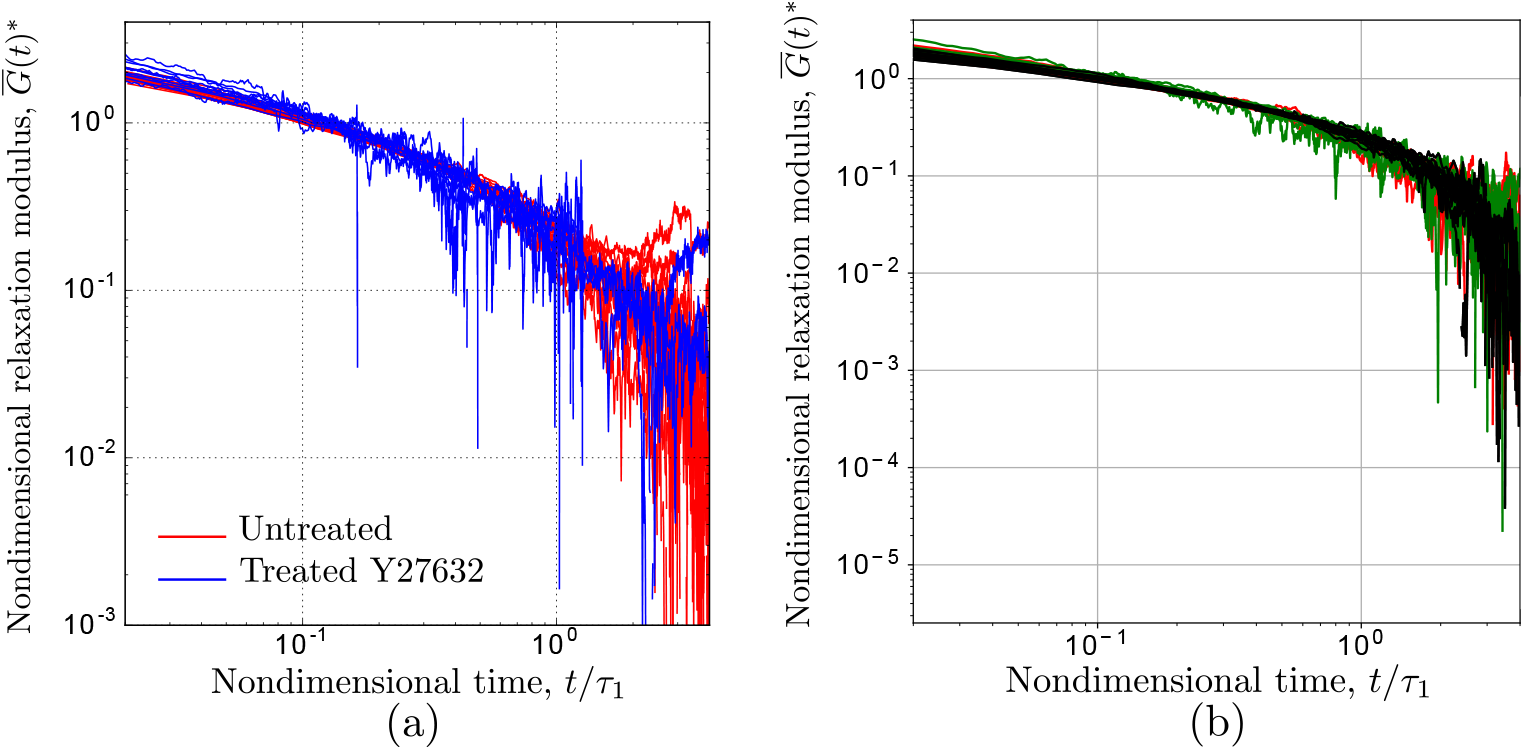
Master curve for the relaxation response. (a) Experimental relaxation curves scaled using the characteristic time *τ*_1_and the effective stiffness of the Fractional Maxwell Model *k*_eff_ The red curves are the experimental data for the untreated monolayers, while the blue curves are the experimental data for the Y27632 treated monolayers. All the data collapse onto a master curve demonstrating that they possess a similar value of *β*. (b) Scaling of the relaxation responses for the other treatments: black DMSO, red CK666, green SMIFH2. All curves collapse to one master curve since they possess a similar value of *β*, the springpot exponent.

### 4 Master curve for the relaxation response

The fact that untreated and treated monolayer can be fitted with the same model, sharing similar values of the exponent *β*, suggests the possibility of identifying a master curve that describes all the experimental data. For this purpose, we simplify the analysis by assuming that the loading is instantaneously applied, thus the stress can be written as *σ*(*t*) = *G*(*t*) *ϵ*_0_, where *G*(*t*) is the relaxation modulus and *ϵ*_0_ the applied stress. By re-writing the relaxation modulus in equation 2 in the main manuscript in terms of the characteristic relaxation time and removing the shift introduced by the addition of a spring in parallel

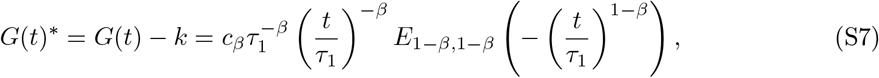

we can identify the additional non-dimensionalising factor for the relaxation modulus *k*_eff_

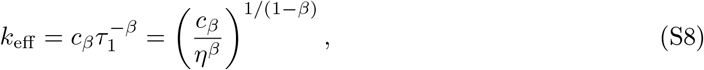

which represents the *effective* stiffness of the Fractional Maxwell Model (FMM), the dissipative branch. Note that *τ*_1_can be also derived from the definition of the characteristic relaxation time for the Maxwell model as *τ*_1_ = *η/k*_eff._ *τ*_1_ is independent from the spring stiffness since the stiffness *k* only shifts the stress towards higher values, as seen in equation 2.

The nondimensionalized relaxation modulus 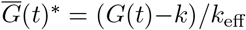, is now a function of only two parameters 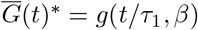, where the time has been nondimensionalized using the characteristic relaxation time *τ*_1_, while the relaxation modulus has been normalized using the effective stiffness *k*_eff_ Consistent with the observation that their short time-scale response is comparable, after rescaling the relaxation curves in figure 2 (b)-(c), all the curves collapse onto one master curve (see figure S4 (a)). From the fitting of the relaxation responses of treated monolayers, we observe that *β* remains constant. Therefore, the nondimensional relaxation modulus can be considered a function of only one parameter 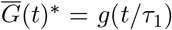, because *β* remains constant. Because of this, all the responses obtained for the other treatments collapse to one master curve (figure S4 (b)).

### 5 Statistical analysis

Statistical analysis of the data was performed in *Julia* language using a Wilcoxon rank-sum test. Changes with statistical power greater than 0.01 were considered non-significant (ns). Dataset different *p*< 0.01 are denoted by an asterisk (*).

**Figure S5:**
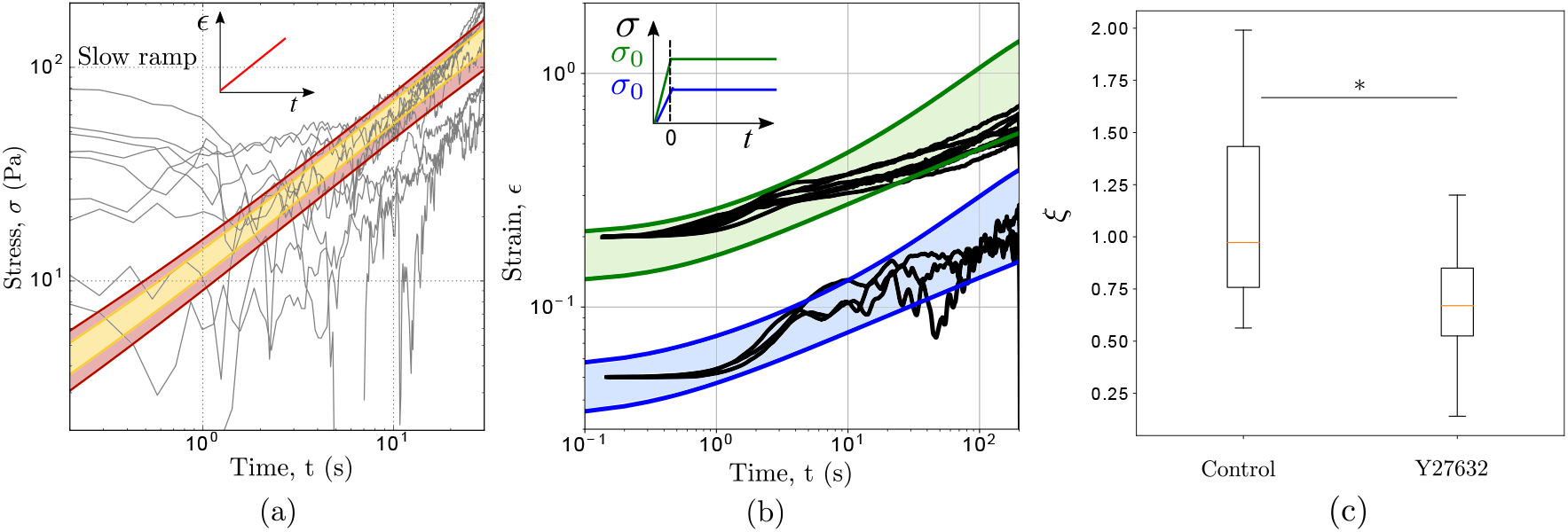
(a) Predicted stress response of the Y27632 treated monolayers when subjected to a slow stretch (1%/s) using the mechanical parametrization determined from stress relaxation experiments. The predicted responses (95% confidence interval red areas and 70% yellow areas) are in good agreement with the experimental data (black curves). (b) The creep response of epithelial monolayers in which actomyosin contractility is inhibited. The creep is performed at 60 Pa (blue area) and 230 Pa (green area). The black lines are the experimental results. (c) *ξ* value for untreated and treated monolayers which give raise to different qualitative behaviours (*p* < 0.01).

The material parameters measured from the relaxation experiment show some variability. We assume that they follow a Gaussian distribution. To take this into account, we take an interval for each of the parameters equal to Mean 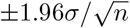, where *σ* here is the standard deviation and *n* the number of samples. The colored areas within the two curves obtained by taking lower and upper bounds of the parameters is where the predicted response of the material will lay.

### 6 Mechanical testing of single epithelial cells

Experiments were performed on MDCK II epithelial cells previously trypsinised and plated in a Petri dish and left to settle for 10-30 minutes. Before cells started to spread, they were tested while still in rounded shape.

Relaxation experiments were performed as described in [6]. A CellHesion 200 Atomic Force Microscope was mounted on a scanning laser confocal microscope. Tipless silicon SPM-Sensor cantilevers with nominal spring constant of 0.03N m^−1^ were used to perform the experiments. The cantilever spring constant for each experiment was calibrated using the thermal noise fluctuation method and it ranged between 0.055-0.06 N m^−1^.

The target force required to indent the cell by ∼ 30% was estimated by approximating the cell height. The latter was given by the difference between the force-displacement curves acquired on the cell and a glass region close to it. Cells were subjected to the target force of 5-40 nN at a rate of 75% s^−1^. The cantilever beam was maintained at constant height for 150 s while the evolution of the force was recorded.

### 7 The complex modulus of the novel generalized fractional model

Although the relaxation modulus *G*(*t*) and the creep compliance *J* (*t*) each fully summarize the viscoelastic linear response of a material, the time-dependent response is often characterized using the complex elastic modulus, which captures the ratio and phase difference between stress and strain under oscillatory deformation, which is particularly relevant to biomaterials because these are often subjected to cyclical stresses. The complex modulus can be decomposed into a real *G*′ (*ω*) and an imaginary *G*″ (*ω*) part, which represent respectively the storage modulus that accounts for the elastic response, and the loss modulus that accounts for the dissipative response. The complex modulus of the viscoelastic model presented here can be derived from the Laplace transform of the creep compliance *J* (*s*) and is given by:

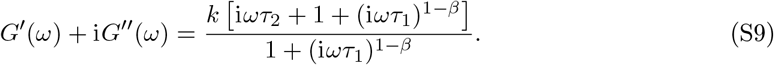

By evaluating the real and imaginary parts of the equation above, we find that the storage and loss moduli are respectively given by

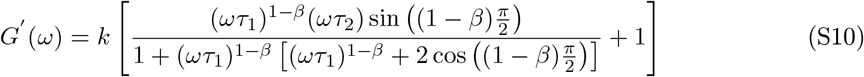

and

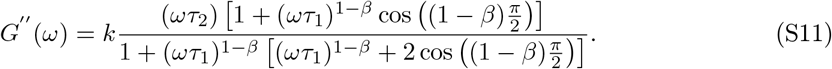

Storage and loss moduli predicted for the monolayers using the parameters obtained from the relaxation experiments both show a power-law behaviour at high frequencies with the same exponent, as shown in figure S10. By contrast, at low frequencies the material exhibits solid-like behaviour [13], [14]. Two special cases can be distinguished at the upper and lower limit of *β*; when the two moduli reduce respectively to those of the Standard Linear Solid model and the Kelvin-Voigt model, as confirmed in figure S10 (b). By letting *β* vary between 0 and 1, we can observe transitory behaviours between an SLS and a Kelvin-Voigt model (see figure S11).

**Figure S6:**
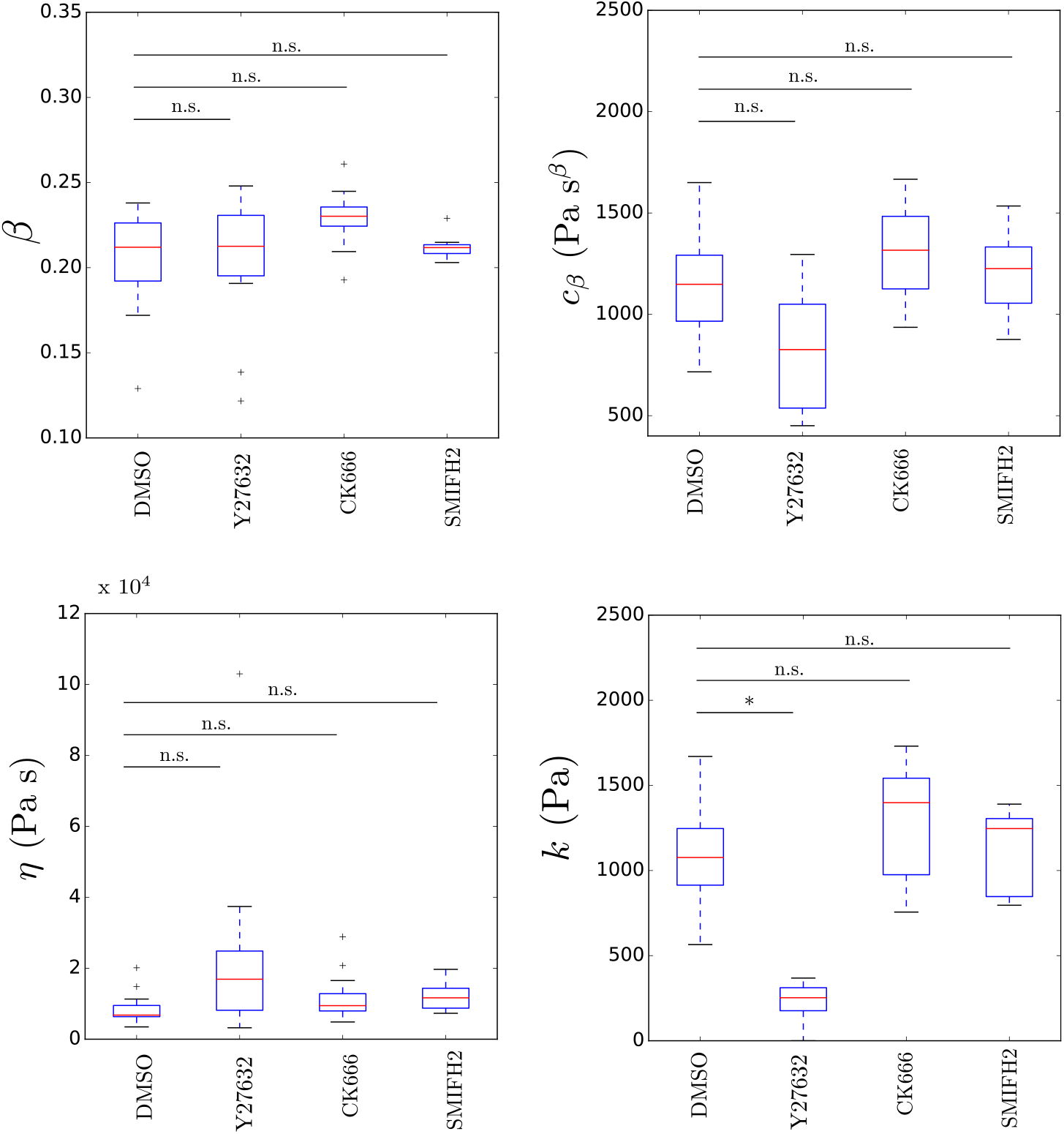
Boxplots comparing the model parameters of monolayers subjected to chemical treatments targeting the actomyosin cytoskeleton. The analysis of the relaxation response of monolayers treated with CK666 that prevents actin polymerisation through Arp2/3 and those treated with SMIFH2 that prevents formins-mediated actin polymerisation revealed that formin inhibition increases the relaxation characteristic time. We can attribute such increment to a higher viscosity *η*, whilst all the other parameters remain almost unchanged. (*β*: *p* = 0.64 for Y27632, *p* = 0.019 for CK666, *p* = 0.98 for SMIFH2, *c*_*β*_: *p* = 0.015 for Y27632, *p* = 0.053 for CK666, *p* = 0.50 for SMIFH2, *η*: *p* = 0.04 for Y27632, *p* = 0.053 for CK666, *p* = 0.022 for SMIFH2, *k*: *p* < 0.01 for Y27632, *p* = 0.19 for CK666, *p* = 0.60 for SMIFH2, for all compared to DMSO.) All drugs were dissolved in Dimethylsulfoxide (DMSO). Therefore, the parameters obtained from the fitting of the relaxation response of the treated monolayers were compared to those obtained from control experiments carried out in the presence of DMSO alone.

**Figure S7:**
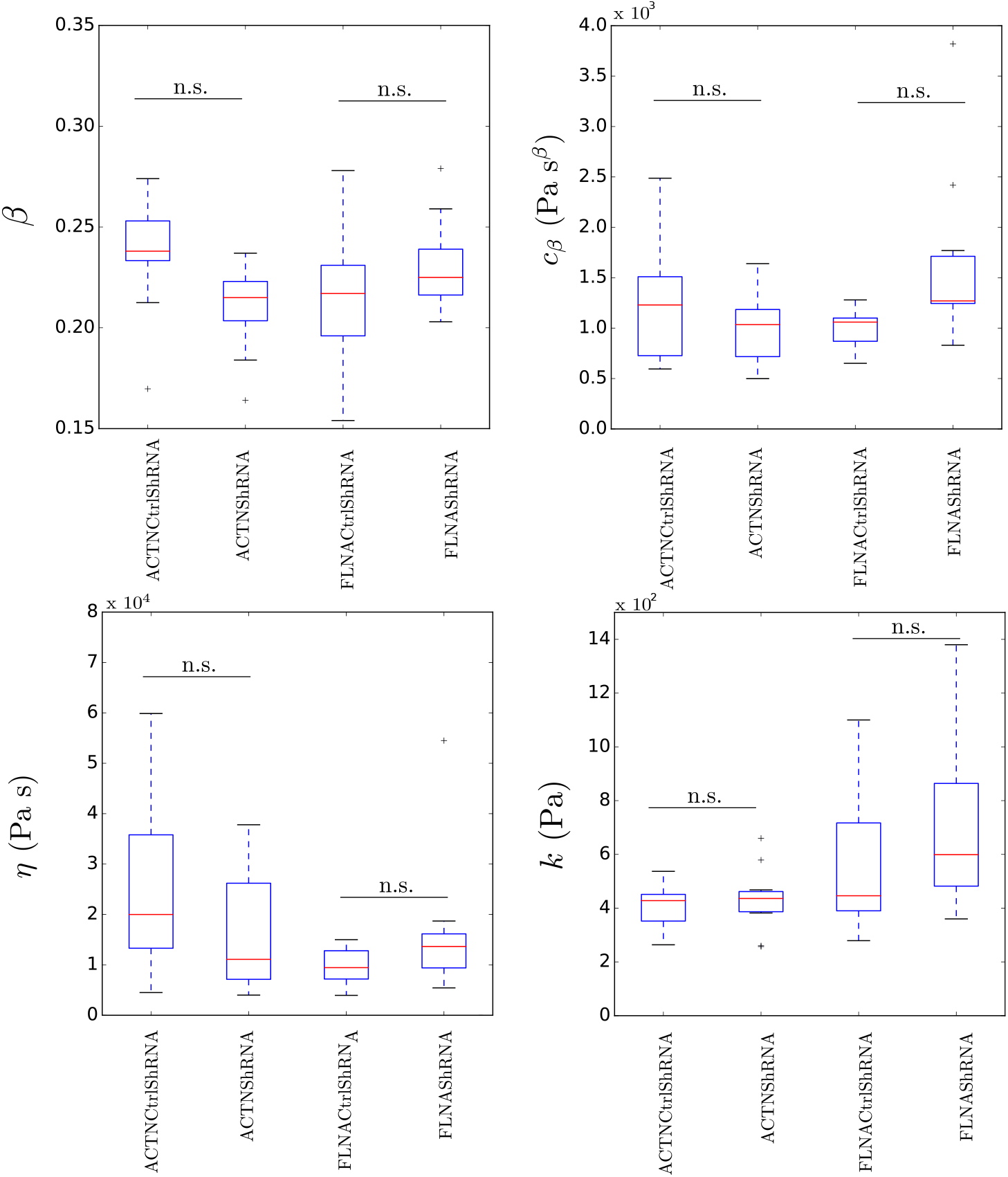
Boxplots comparing the model parameters of monolayers in which the F-actin crosslinkers filamin A (FLNAShRNA) or *α*-actinin 4 (ACTNShRNA), the two most abundant actin crosslinkers identified in previous RNAseq experiments [6], were perturbed. We confirm that crosslinkers do not play a role in stress dissipation following extension (*β*: *p* = 0.03 for ACTNShRNA and *p* = 0.18 for FLNAShRNA, *c*_*β*_: *p* = 0.71 for ACTNShRNA and *p* = 0.018 for FLNAShRNA, *η*: *p* = 0.28 for ACTNShRNA and *p* = 0.09 for FLNAShRNA, *k*: *p* = 0.78 for ACTNShRNA and *p* = 0.25 for FLNAShRNA, compared to their respective controls, ACTNCtrlShRNA and FLNACtrlShRNA.

**Figure S8:**
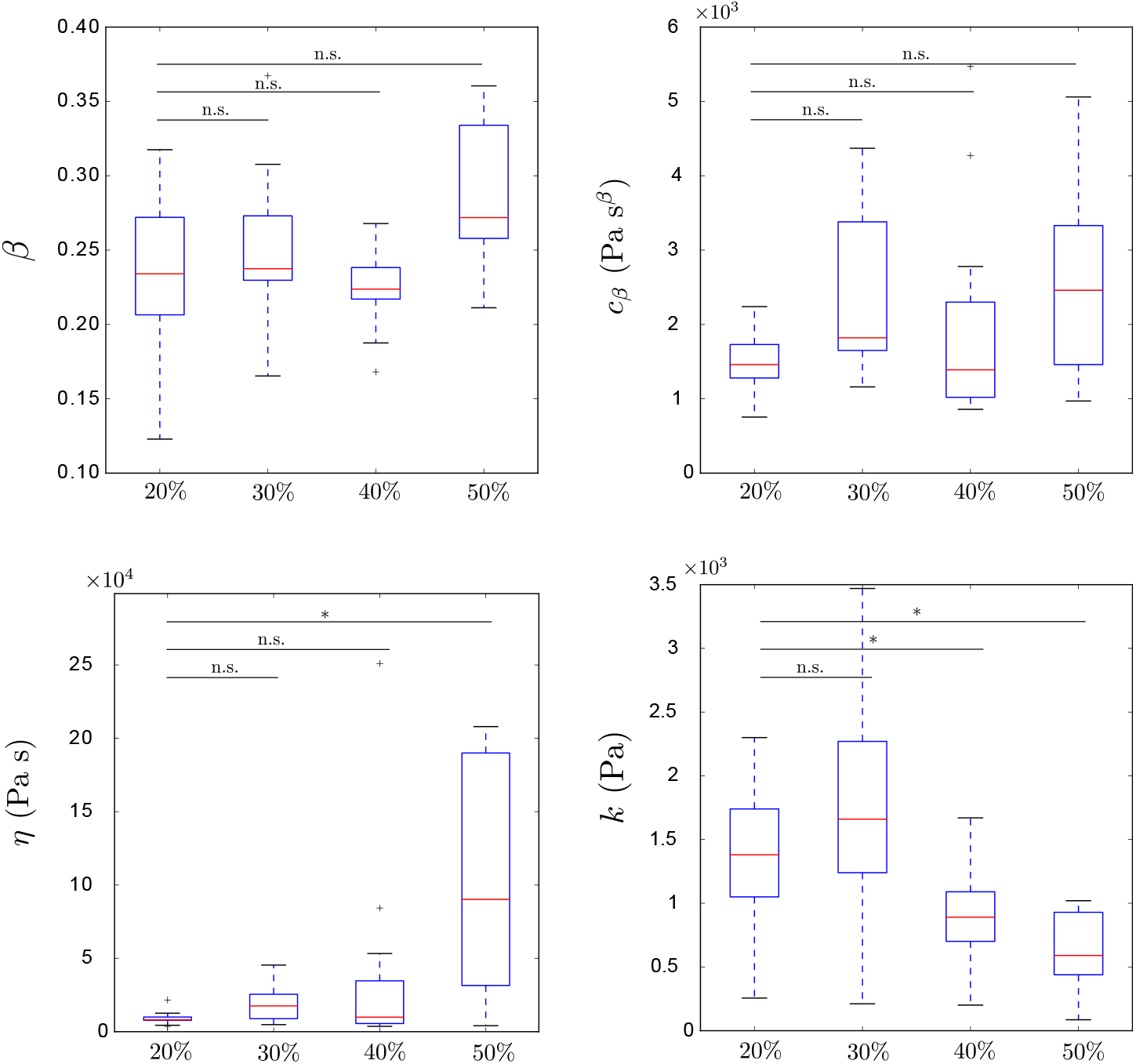
Test of linearity. The average parameters obtained from the fitting of the relaxation curves for tissues subjected to different strain amplitudes (samples’ number for each value of strain is 13): (a) power-law exponent *β* (*p* = 0.51 for 30%, *p* = 0.51 for 40%, *p* = 0.03 for 50%). (b) spring-pot coefficient *c*_*β*_ (*p* = 0.03 for 30%, *p* = 0.90 for 40%, *p* = 0.04 for 50%) (c) viscosity *η* (*p* = 0.05 for 30%, *p* = 0.29 for 40%, *p* = 0.0012 for 50%) (d) stiffness *k* (*p* = 0.44 for 30%, *p* = 0.007 for 40%, *p* < 0.001 for 50%).

**Figure S9:**
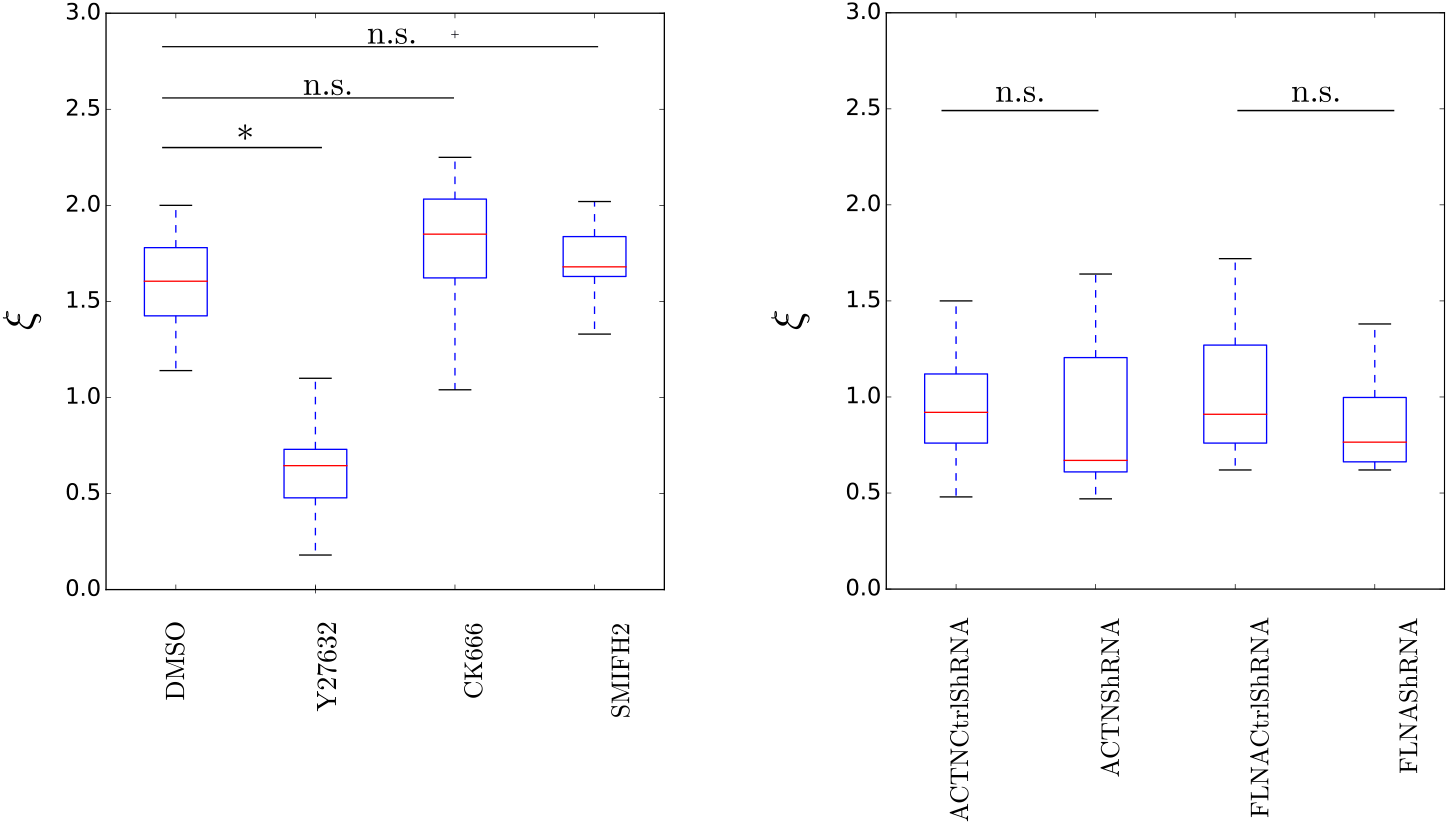
Boxplots of the ratio of the two characteristic times for monolayers (a) in which the actin network is perturbed. (*p* < 0.01 for Y27632, *p* = 0.07 for CK66, *p* = 0.37 for SMIFH2, all compared to DMSO (b) Monolayers in which the crosslinkers are perturbed (*p* = 0.68 for ACTNShRNA and *p* = 0.34 for FLNAShRNA both compared to their controls). A slight increase is observed when treated with SMIFH2, whilst no effect is observed when crosslinking is perturbed.

**Table S2:** Fitted parameters.

| Sample | $\eta$ (Pa s) | $\beta$ | $c_\beta$ (Pa s$^\beta$) | k (Pa) | $\tau_1$ (s) | $\tau_2$ (s) | $\xi$ |
| --- | --- | --- | --- | --- | --- | --- | --- |
| Epithelial MDCK monolayer [6] | $1 \times 10^4 \pm 8 \times 10^3$ | $0.22 \pm 0.02$ | $1360 \pm 420$ | $760 \pm 340$ | 13 | 13 | 1 |
| Epithelial MDCK monolayer, Y27632 treated [6] | $5 \times 10^4 \pm 2 \times 10^4$ | $0.28 \pm 0.05$ | $1120 \pm 441$ | $187 \pm 100$ | 195 | 270 | 0.7 |
| Single epithelial MDCK cell [6] | $870 \pm 720$ | $0.14 \pm 0.03$ | $109 \pm 20$ | $203 \pm 32$ | 10 | 4 | 2.6 |
| Single articular chondrocyte [7] | ∞ | 0.70 | 140 | 133 | ∞ | ∞ | ∞ |
| Collagen fibrils [8] | $722$ (MPa s$^\beta$) | 0.14 | $14$ (MPa s$^\beta$) | $83$ (MPa) | 100 | 10 | 11 |
| Single C2-7 myogenic cells [9] | $4 \times 10^5$ | 0.26 | 885 | 0.0 | ∞ | - | - |
| Cytoplasm [10] | 35 | 0.3 | 0.7 | 0.16 | 270 | 218 | 1 |
| Yolk cell [10] | $16 \times 10^3$ | 0.15 | 0.62 | 0.52 | ∞ | - | - |
| HeLa Kyoto cell cortex [11] | $170$ (mN/m s) | 0.22 | $48$ (mN/m s$^\beta$) | $5$ (mN/m) | - | - | - |
| Human primary immune cells [12]. | ∞ | 0.26 | 600 | 0.0 | - | - | - |

**Figure S10:**
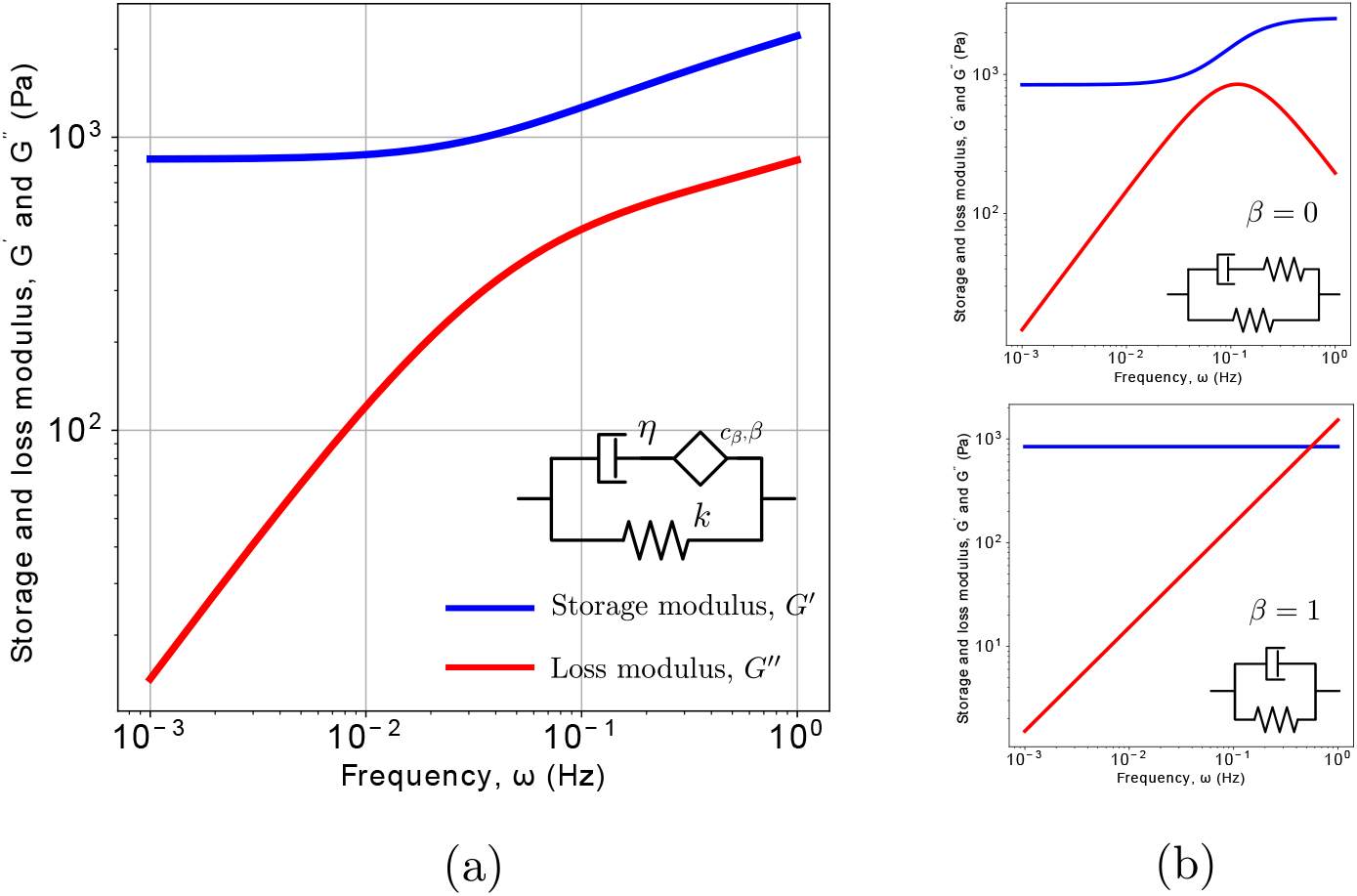
Storage and loss moduli of the fractional model shown in figure 2 (a). (a) The blue and red lines are respectively the storage and loss modulus of the viscoelastic model introduced here using the material parameters extracted from the relaxation experiment for epithelial monolayers shown on figure 1 (a) (*β* = 0.22, *c*_*β*_ = 1.3 10_3_Pa s^*β*^, *k* = 760 Pa, *η* = 1.4 10_4_ Pa s, *τ*_1_*/τ*_2_ = 1). Moduli for two extreme cases. When *β* = 0 the model behaves as a Standard Linear Solid model (top), when *β* = 1 the model behaves as a Kelvin mode (bottom). The other parameters have been kept constant.

**Figure S11:**
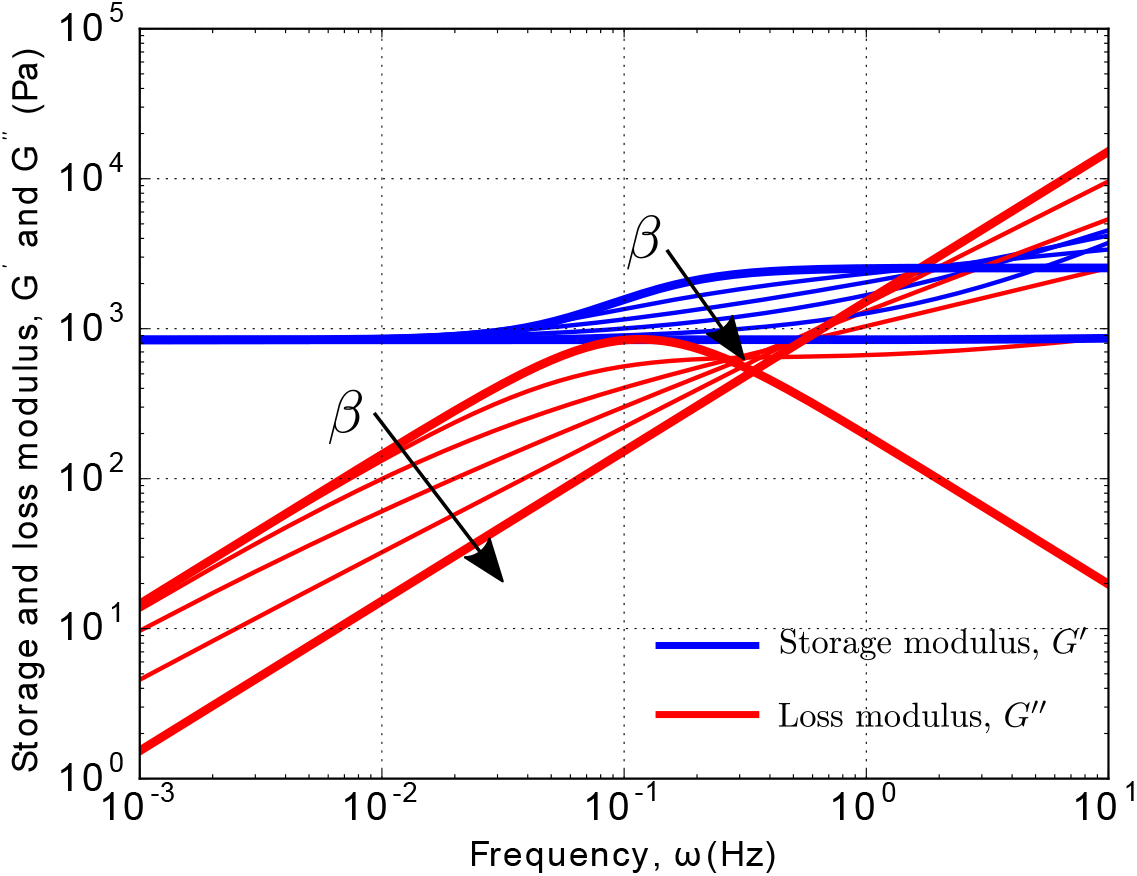
Intermediate behaviours from Kelvin model to Standard Linear Solid Model (*cβ* = 1.3 × 10^3^ Pa s^*β*^, *k* = 800 Pa, *η* = 1 × 10_4_ Pa s, *β* = 0, 0.2, 0.4, 0.6, 0.8, 1. The blue and red lines are respectively the storage and loss moduli. Note that by changing the value of *β* the ratio between the two characteristic time scales changes and it is respectively given by *τ*_1_*/τ*_2_ = 0.46, 0.85, 2.08, 12.5, 270, ∞. The thicker curves are those presented in figure S10 in the main text.

**Figure S12:**
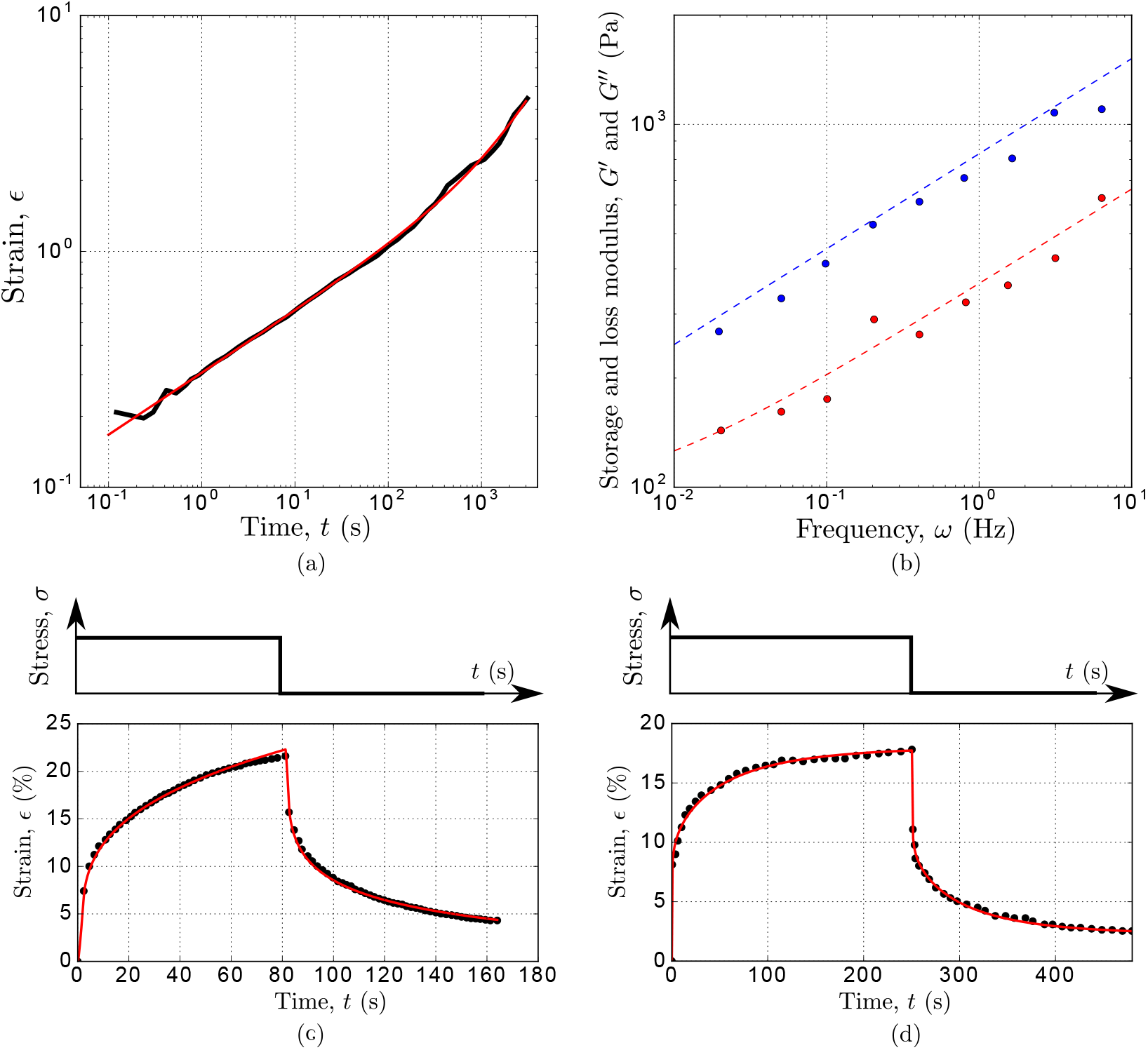
Examples of the application of the fractional viscoelastic model for the analysis of the mechanical response in biology. (a) Fitting of the creep response of C2-7 myogenic cells derived from skeletal muscle of adult CH3 mice (original data from [9]) with a special case of the model presented here. Fitted parameters: *η* = 4 ×10^5^Pa s, *c*_*β*_ = 885 Pa s^*β*^, *β* = 0.26 (b) The prediction of the storage and loss modulus (respectively blue and red dashed lines) of C2-7 cells using the parameters obtained from the fitting of the creep response in (a) are in good agreement with the experimental data obtained from [15] (c) Fitting of the creep and recovery response (red line) of the cytoplasm of the blastomere, and (d) the yolk cell (original data from [10], black dots) using the viscoelastic model presented here. Fitted parameters for cytoplasm: *η* = 35 Pa s, *c*_*β*_ = 0.7 Pa s^*β*^, *β* = 0.3, *k* = 0.156 Pa. Fitted parameters yolk cell: *η* = 16 ×10_3_ Pa s, *c*_*β*_ = 0.62 Pa s^*β*^, *β* = 0.15, *k* = 0.52 Pa.

**Figure S13:**
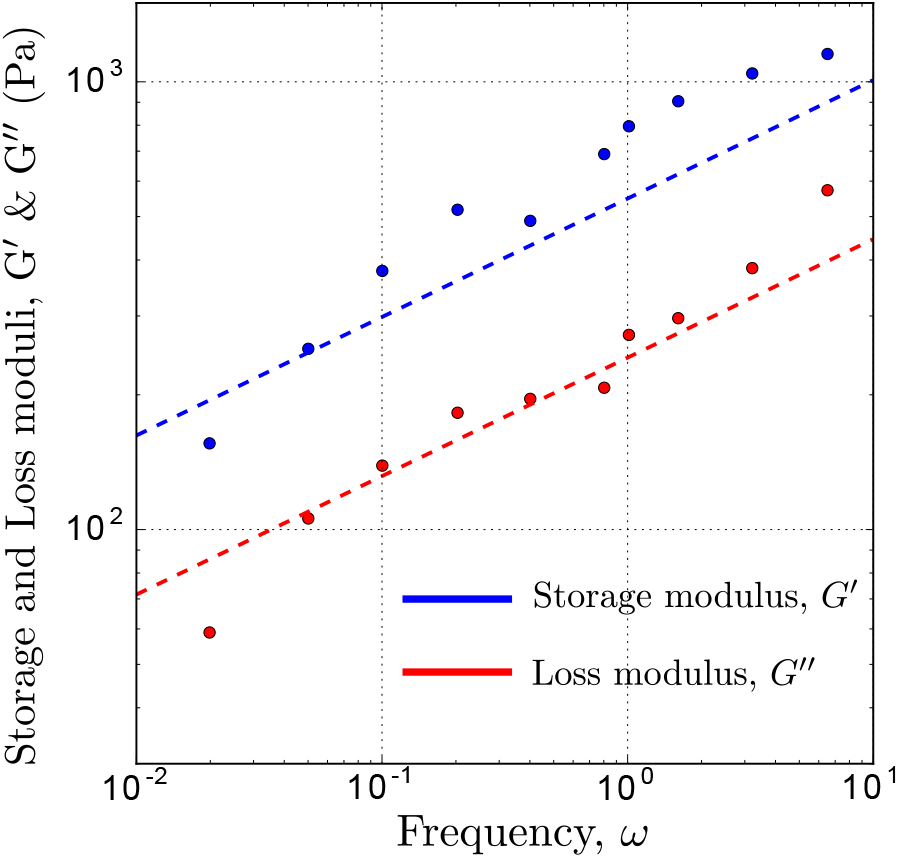
Fitting of storage and loss moduli from a single cell dynamic test of human primary immune cells. Original data from [12]. Fitted parameters: *η* = Pa s, *cβ* = 600 Pa s*β*, *β* = 0.26, k = 0. Note that the high viscosity and the zero value of the spring are directly obtained from the fitting, without constrains on the parameters.

